# Overcoming antibody-resistant SARS-CoV-2 variants with bispecific antibodies constructed using non-neutralizing antibodies

**DOI:** 10.1101/2023.10.26.564289

**Authors:** Tetsuya Inoue, Yuichiro Yamamoto, Kaoru Sato, Yuko Nakamura, Yoshimi Shimizu, Motohiko Ogawa, Taishi Onodera, Yoshimasa Takahashi, Takaji Wakita, Mika K. Kaneko, Masayoshi Fukasawa, Yukinari Kato, Kohji Noguchi

**Affiliations:** Laboratory of Molecular Targeted Therapy, Faculty of Pharmaceutical Sciences, Tokyo University of Science, Yamazaki 2641, Noda, Chiba 278-8510, Japan; Department of Biochemistry and Cell Biology, National Institute of Infectious Diseases, 1-23-1, Toyama, Shinjuku-ku, Tokyo 162-8640, Japan; Department of Pharmaceutical Sciences, Teikyo Heisei University, 4-21-2 Nakano, Nakano-ku 164-8530, Japan; Department of Virology I, National Institute of Infectious Diseases, 1-23-1, Toyama, Shinjuku-ku, Tokyo 162-8640, Japan; Research Center for Drug and Vaccine Development, National Institute of Infectious Diseases, 1-23-1, Toyama, Shinjuku-ku, Tokyo 162-8640, Japan; National Institute of Infectious Diseases, 1-23-1, Toyama, Shinjuku-ku, Tokyo 162-8640, Japan; Department of Molecular Pharmacology, Tohoku University Graduate School of Medicine, 2-1 Seiryo-machi, Sendai, Miyagi 980-8575, Japan; Department of Antibody Drug Development, Tohoku University Graduate School of Medicine, 2-1 Seiryo-machi, Sendai, Miyagi 980-8575, Japan

## Abstract

A current challenge is the emergence of SARS-CoV-2 variants, such as BQ.1.1 and XBB.1.5, that can evade immune defenses, thereby limiting antibody drug effectiveness. Emergency-use antibody drugs, including the widely effective bebtelovimab, are losing their benefits. One potential approach to address this issue are bispecific antibodies which combine the targeting abilities of two antibodies with distinct epitopes. We engineered neutralizing bispecific antibodies in the IgG-scFv format from two initially non-neutralizing antibodies, CvMab-6 (which binds to the receptor-binding domain [RBD]) and CvMab-62 (targeting a spike protein S2 subunit epitope adjacent to the known anti-S2 antibody epitope). Furthermore, we created a bispecific antibody by incorporating the scFv of bebtelovimab with our anti-S2 antibody, demonstrating significant restoration of effectiveness against bebtelovimab-resistant BQ.1.1 variants. This study highlights the potential of neutralizing bispecific antibodies, which combine existing less effective anti-RBD antibodies with anti-S2 antibodies, to revive the effectiveness of antibody therapeutics compromised by immune-evading variants.

## INTRODUCTION

SARS-CoV-2 vaccines, including a new type of mRNA vaccine, have been developed and demonstrated to be highly effective in combating the COVID-19 pandemic. In addition, SARS-CoV-2 RNA polymerase and protease inhibitors have been developed as small-molecule compounds.^1^ Specific monoclonal antibodies with virus-neutralizing activity are another powerful approach for the treatment or prevention of SARS-CoV-2 infection.^2–6^ SARS-CoV-2 antibody therapeutics demonstrating potent virus neutralization activity have been developed using monoclonal antibodies isolated from the B cells of patients with COVID-19.^7–11^ Notably, potent inhibitory antibody therapeutics mostly block the binding between the receptor-binding domain (RBD) of the SARS-CoV-2 spike protein and the cellular receptor angiotensin-converting enzyme 2 (ACE2). These results suggest that the binding interface between ACE2 and the RBD is the optimal target site for potent neutralizing antibodies. Furthermore, neutralizing antibodies that bind to the N-terminal domain or the S2 region of the spike protein have also been identified.^12–16^

Unfortunately, various SARS-CoV-2 variants have acquired immune evasion abilities, even in individuals who have received vaccinations.^17–22^ Interestingly, these variants contain amino acid mutations in the RBD that allow the viruses to escape capture by monoclonal antibodies, resulting in antibody resistance. For example, recent Omicron variants, such as BA.4/5 and BA.2.75, have shown resistance to many monoclonal neutralizing antibodies.^23–28^ Even bebtelovimab,^29^ which remains effective against many variant strains, has seen its effectiveness decreased against the recent variants BQ.1 and XBB.^30, 31^ The direct correlation between mutations in the RBD-binding site and immune evasion suggests that the most efficient mode of action of neutralizing antibodies, which directly target ACE2-RBD binding, may be the most vulnerable to antibody resistance.

Targeting the RBD with a single epitope demonstrating SARS-CoV-2 neutralizing activity has advantages and disadvantages. Antibody resistance issues undermine the value of the monoclonal antibodies that have been developed to date and pose a significant obstacle to the development of new monoclonal antibody therapeutics; thus, strategies to overcome it are highly desired. Therefore, it is important to explore alternative antibody therapeutics that are based on various mechanisms and innovations. Combining multiple antibodies into a cocktail or creating bispecific or multispecific molecules may enhance the efficacy of antibody therapeutics against viral immune evasion.^32, 33^ These approaches mitigate the effects of resistance mutations by targeting multiple viral epitopes.

Exploring non-RBD regions or other sites as broad-spectrum neutralizing epitopes is important in research on antibody therapeutics with novel pharmacological actions. With the development of broad-spectrum antibody therapeutics, neutralizing antibodies that target regions with highly conserved cold spots and non-RBD sites in various SARS-CoV-2 strains have been studied. Antibodies targeting the S2 region of the spike protein, specifically the highly conserved stem helix region, show relatively broad neutralizing activity, although their inhibitory activity is weaker than that of many RBD-binding neutralizing antibodies.^34–41^ To overcome the weaknesses of non-RBD-targeting antibodies, recombinant antibodies can be created by combining them with other molecules to generate powerful inhibitory activity against viral infection. Furthermore, bispecific antibodies that combine with non-neutralizing epitopes are promising therapeutic antibodies.^42, 43^ These strategic concepts have the potential to generate a diverse range of SARS-CoV-2–neutralizing antibodies with various pharmacological mechanisms. This approach is expected to be effective in combating the emergence of immune evasion mutations.

In this study, we developed bispecific neutralizing antibodies by combining two types of antibodies that do not directly inhibit the binding of the spike protein to ACE2: one antibody binds to the RBD but lacks neutralizing activity, and the other targets a highly conserved epitope in the S2 region with weak neutralizing activity. This approach led to the discovery of a clone with enhanced inhibitory effects and broad-spectrum infection-blocking activity. Moreover, we combined the single-chain variable fragment (scFv) of bebtelovimab, which is a therapeutic anti-RBD antibody^29^ that had become ineffective owing to the emergence of the resistant variants BQ.1 and XBB,^30, 31^ with our anti-S2 antibody CvMab-62. We found that this bispecific antibody overcame the resistance of BQ.1.1 to bebtelovimab. Thus, bispecific antibodies combining the S2 antibody with other RBD-targeting antibodies may be a promising and optional module to restore efficacy against antibody-resistant SARS-CoV-2 variants.

## RESULTS

### Bispecific antibodies were generated from the combination of the non-neutralizing anti-RBD antibody CvMab-6 and anti-S2 antibody CvMab-62

It has been reported that bispecific antibodies, combined with non-neutralizing antibodies that recognize highly conserved regions, demonstrate broad neutralizing activity.^42^ Furthermore, bispecific antibodies that recognize both the RBD and S2 regions have been reported to exhibit better neutralizing activity than their parental monoclonal antibodies.^44^ Therefore, we generated anti-SARS-CoV-2 bispecific antibodies from non-neutralizing anti-RBD and anti-S2 antibodies.

CvMab-6 targeting RBD and CvMab-62 targeting S2 were broadly reactive antibodies, and their binding to spike proteins of variants including the Wuhan strain, D614G, Alpha, Delta, and Omicron BA.1, were confirmed by western blot and indirect immunofluorescence analyses (Figures 1A, 1B, 1C, and 1D, respectively). The anti-S2 antibody, CvMab-62, but not the anti-RBD antibody, CvMab-6, showed weak but selective inhibition against pseudotyped and authentic SARS-CoV-2 infections only at high concentrations (Figures 1E–1G). The combination of CvMab-6 and CvMab-62 showed no synergistic effects (Figure 1H). These results indicated that the anti-S2 antibodies CvMab-62 and anti-RBD CvMab-6 are non-neutralizing antibodies compared to the previously reported neutralizing antibodies.

**Figure 1.**
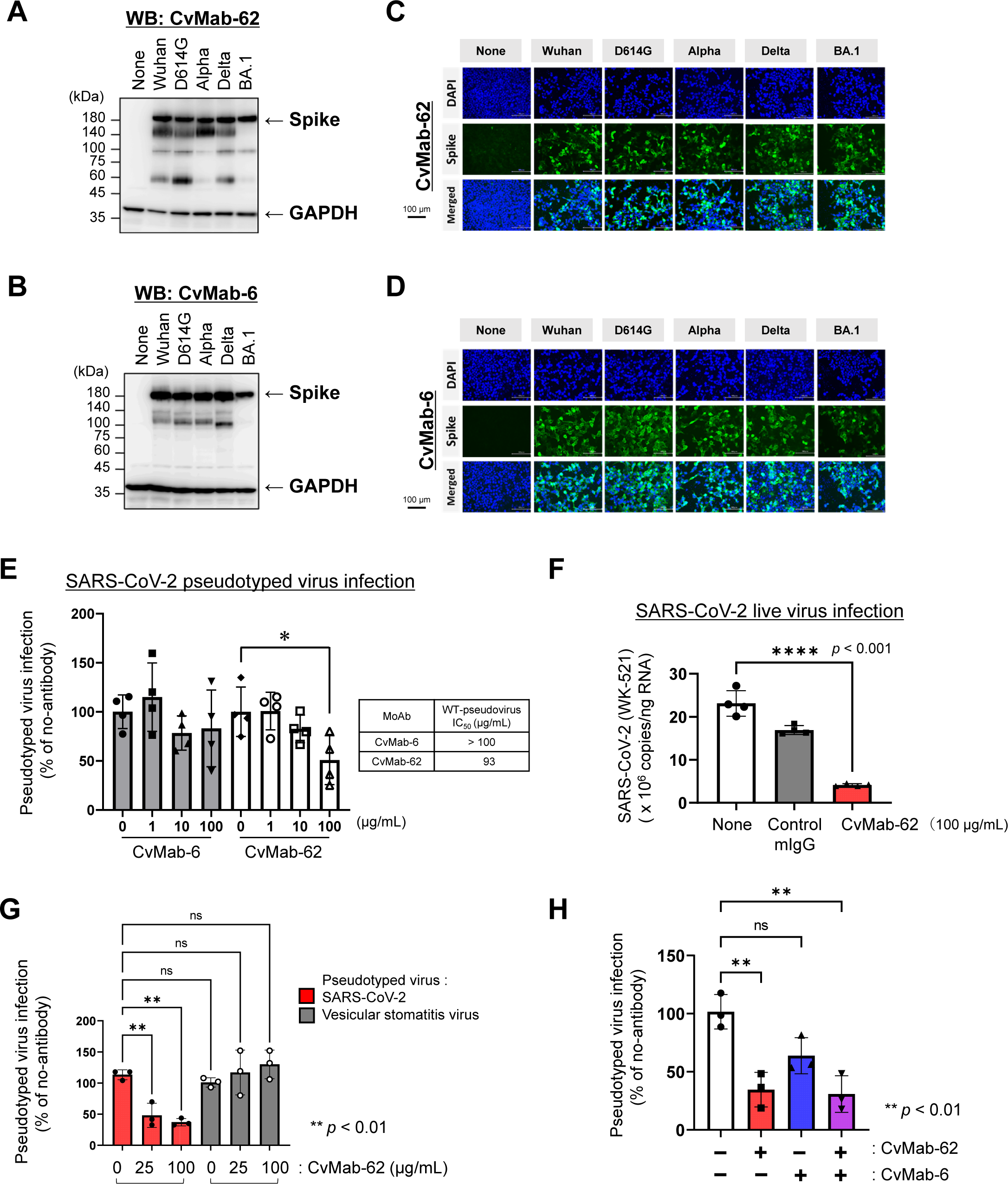
Broad reactivities of anti-S2 CvMab-62 and anti-RBD CvMab-6 antibodies. (A) Western blot analysis using CvMab-62. Spike proteins of Whuan-Hu-1, D614G, Alpha, Delta, and Omicron BA.1 expressed in HEK293T cells detected using anti-S2 CvMab-62 antibody. (B) Western blotting analysis of CvMab-6. Spike proteins of Whuan-Hu-1, D614G, Alpha, Delta, and Omicron BA.1 expressed in HEK293T cells detected using anti-RBD CvMab-6 antibody. (C) Indirect immunofluorescence analysis of CvMab-62. Spike proteins of Whuan-Hu-1, D614G, Alpha, Delta, and Omicron BA.1 expressed in HEK293T cells, probed with anti-S2 CvMab-62 antibody, and visualized using secondary anti-mouse IgG-Alexa488 (green signals). Nuclei are counter-stained with DAPI. (D) Indirect immunofluorescence analysis using CvMab-6. Spike proteins of Whuan-Hu-1, D614G, Alpha, Delta, and omicron BA.1 expressed in HEK293T cells, probed with anti-RBD CvMab-6 antibody, and visualized using secondary anti-mouse IgG-Alexa488 (green signals). Nuclei are counter-stained with DAPI. (E) Ineffectiveness of CvMab-6 and CvMab-62 against Wuhan-Hu-1 type SARS-CoV-2 pseudotyped virus. A pseudotyped virus with a luciferase reporter gene was preincubated with antibodies and used to infect VeorE6/TMPRSS2 cells. The infection ratio was evaluated by measuring cellular luciferase activity 3 days post-infection (n=4). (F) Low effectiveness of CvMab-62 against Wuhan-Hu-1 type authentic live SARS-CoV-2 virus. Live SARS-CoV-2 (WK-521) was preincubated with CvMab-62 or control mouse IgG antibodies and used to infect VeorE6/TMPRSS2 cells. The infection ratio was evaluated by measuring cellular viral genomic RNA at 1 day post-infection using quantitative PCR (n=4). (G) Selectivity of CvMab-62 against Wuhan-Hu-1 type pseudotyped virus. Wuhan-type SARS-CoV-2 or control VSV pseudotyped viruses were preincubated with the CvMab-62 antibody and used to infect VeorE6/TMPRSS2 cells. Infection ratio was evaluated by measuring cellular luciferase activity at 3 days post-infection (n=3). (H) No synergy was observed between the CvMab-6 and CvMab-62. Wuhan-type SARS-CoV-2 pseudotyped virus with a luciferase reporter gene was preincubated with either each antibody or a cocktail of antibodies, and VeorE6/TMPRSS2 cells were infected. The infection ratio was evaluated by measuring cellular luciferase activity at 3 days post-infection (n=3). Statistical differences were determined using a one-way ANOVA, and a *p*-value < 0.05 was considered statistically significant, \**p* < 0.05, \*\**p* < 0.01, \*\*\**p* < 0.001.

The CvMab-6 epitope corresponds to amino acids 459–478 of the spike protein, determined by an enzyme-linked immunosorbent assay (ELISA) using a synthetic series of SARS-CoV-2-S1-RBD peptides, as shown in Table S1. The amino acid sequence at 459–478 is highly conserved in SARS-CoV-2 and bat RaTG13 spike proteins but not in the spike protein of another bat species, Khosta-2 (Figure 2A). Consistently, CvMab-6 recognized the bat coronavirus RaTG13 spike protein but did not react with the bat coronavirus Khosta2 spike protein (Figures 2B and 2C, respectively). This region was not the binding interface between ACE2 and RBD (Figure 2D), and consistently CvMab-6 did not inhibit ACE2-RBD binding in ELISAs (Figure 2E), confirming that CvMab-6 is a non-neutralizing antibody.

**Figure 2.**
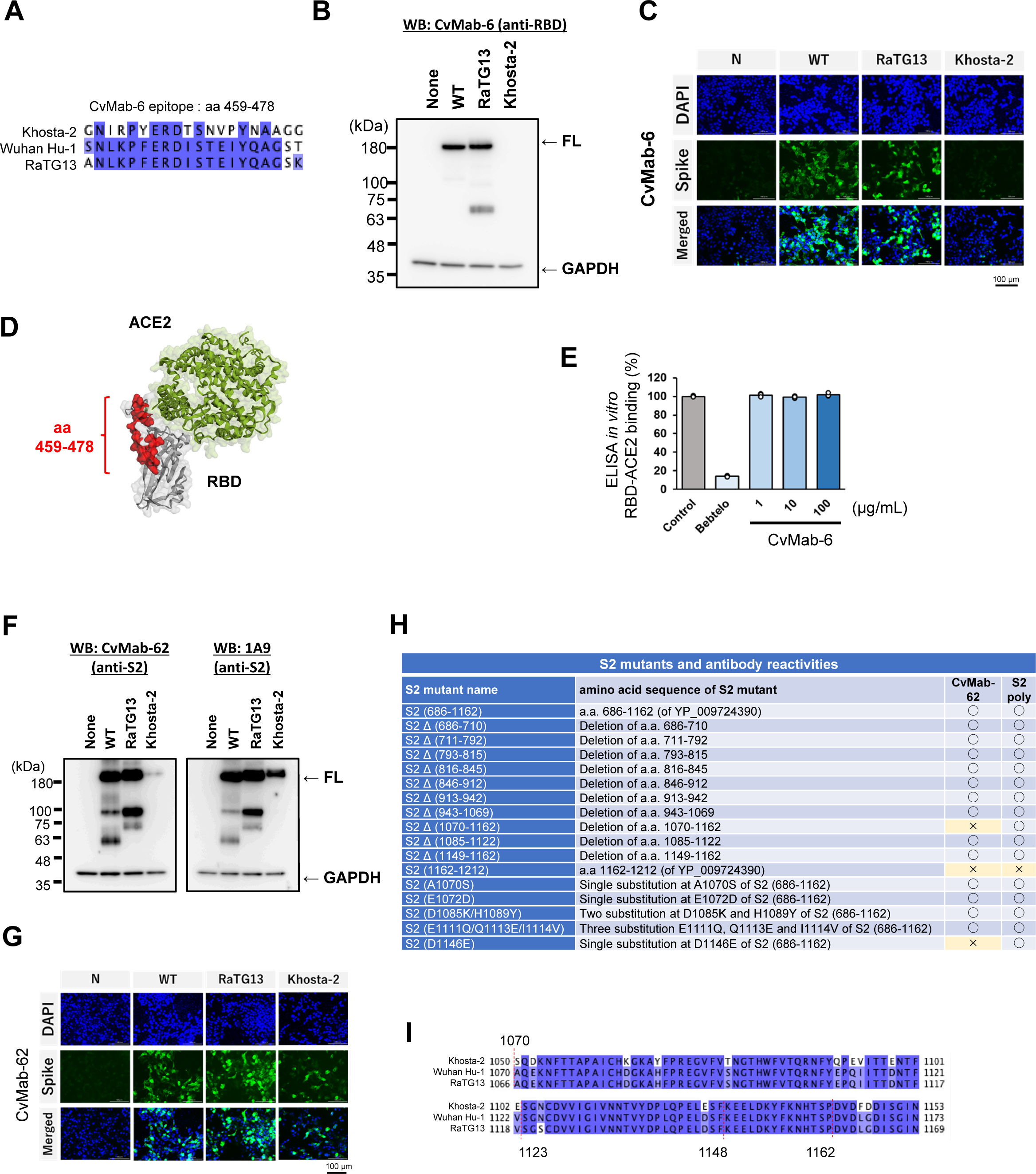
Anti-RBD CvMab-6 and anti-S2 CvMab-62 epitopes. (A) Amino acid alignment of CvMab-6 target element. Amino acid residues corresponding to position 459–478 of the Wuhan-Hu-1 spike protein are aligned with those of bat coronaviruses RaTG13 and Khosta2. (B) Western blot analysis of CvMab-6 on bat coronavirus spike proteins. CvMab-6 did not react with the Khosta2 spike protein. (C) Indirect immunofluorescence analysis of CvMab-6 on bat coronavirus spike proteins. CvMab-6 did not react with the Khosta2 spike protein expressed in HEK293 cells. (D) Structural model of the CvMab-6 target element. The red region represents the CvMab-6 binding site. (E) CvMab-6 did not inhibit RBD-ACE2 binding. Preincubation of RBD with bebtelovimab, but not with CvMab-6, inhibited *in vitro* binding between RBD and ACE2 using ELISA (n=3). (F) Western blot analysis of CvMab-62 on bat coronavirus spike proteins. The reactivity of CvMab-62 with the Khosta2 spike protein was strongly reduced compared to the anti-S2 1A9 antibody. (G) Indirect immunofluorescence analysis showed reduced reactivity of CvMab-62 on bat coronavirus Kohosta2 spike proteins expressed in HEK293T cells. (H) Summary of western blot analysis of the deletion mutant S2 proteins using CvMab-62. CvMab-62 detected a deletion mutant S2 protein lacking amino acid residues 1149–1162, but did not react with the D1146E mutant S2 protein.

The CvMab-62, an anti-S2 antibody, also showed weak reactivity towards Khosta-2, unlike the control anti-S2 antibody 1A9 which recognizes the highly conserved region, Wuhan-Hu-1 spike protein at amino acids position 1029–1192 (Figures 2F and 2G). As summarized in Figures 2H and S1, western blot analysis using the deletion mutants of the S2 protein revealed that CvMab-62 did not interact with the mutants lacking residues 1070–1162. The spike protein of bat Khosta2 differs from SARS-CoV-2 in the corresponding 1070–1162 residue region due to amino acid substitutions such as A1070S, E1072D, D1084K, H1088Y, S1097T, E1111Q, Q1113E, I1114V, D1118E, V1122E, and D1146E (Figure 2I), and additional western blot analyses revealed that mutations A1070S, E1072D, D1084K/H1088Y, QPEV (E1111Q/Q1113E/I1114V), the deletion from 1085–1122, and the deletion from 1149–1162 do not impact CvMab-62 binding, but D1146E mutation disrupts CvMab-62 binding (Figure S1). In addition, the binding of CvMab-62 to the S2 region does not require the presence of residues 1149–1162 (KEELDKYFKNHTSP), a common epitope for most anti-S2 neutralizing antibodies,^34, 45^ indicating that the CvMab-62 epitope is novel within the region of residues 1123–1148, with a particular focus on D1146.

Four IgG-type bispecific antibodies were tested using the original CvMab-6 and CvMab-62 antibodies. Bis1 was generated by fusing the scFv of CvMab-62, which recognizes S2, to the C-terminus of the heavy chain of cCvMab-6, whereas Bis2 was generated by fusing it to the C-terminus of the light chain of cCvMab-6. Bis3 was generated by fusing the scFv of cCvMab-6, which recognizes the RBD, to the C-terminus of the heavy chain of CvMab-62, and Bis4 was generated by fusing it to the C-terminus of the light chain of CvMab-62 (Figure 3A). Purified recombinant antibodies were confirmed using SDS-PAGE and Coomassie brilliant blue staining (Figure 3B). The dual-binding activity of these bispecific antibodies was confirmed by ELISA (Figure S2), and the binding affinities of each recombinant antibody towards the trimeric ectodomain of the spike protein (Figure 3C) and RBD alone (Figure 3D) were evaluated by ELISA. The binding affinity of Bis1 was lower than that of the parental antibodies CvMab-6 and CvMab-62, whereas the binding affinity of Bis2 was comparable to that of the anti-RBD antibody CvMab-6 (Figure 3E). In contrast, Bis3 and Bis4 exhibited binding affinities to the trimeric ectodomain of the Wuhan-type spike protein at a level similar to that of the anti-S2 antibody CvMab-62, and their binding affinities to the RBD were comparable to that of CvMab-6. As expected, these bispecific antibodies did not inhibit *in vitro* binding between the RBD and ACE2 (Figure S3). However, when the antiviral activity of these bispecific antibodies was examined in a pseudotyped virus assay, Bis3 showed the strongest inhibitory activity against the Wuhan, Alpha, Delta, and BA.1 variants (Figure 3F). The IC_50_ value of Bis3 was lower than that of the parental CvMab-62, indicating improved neutralizing activity through bispecific antibody formation. Furthermore, we evaluated the inhibitory activity against SARS-CoV-2 live virus infection, and similarly, Bis3 exhibited the strongest inhibition against the Wuhan, Alpha, Delta, and BA.1 strains among the four types of bispecific antibodies (Figure 3G). These results demonstrate that bispecific antibodies combining non-neutralizing antibodies, particularly those based on S2 antibodies, such as Bis3, can generate antiviral activity.

**Figure 3.**
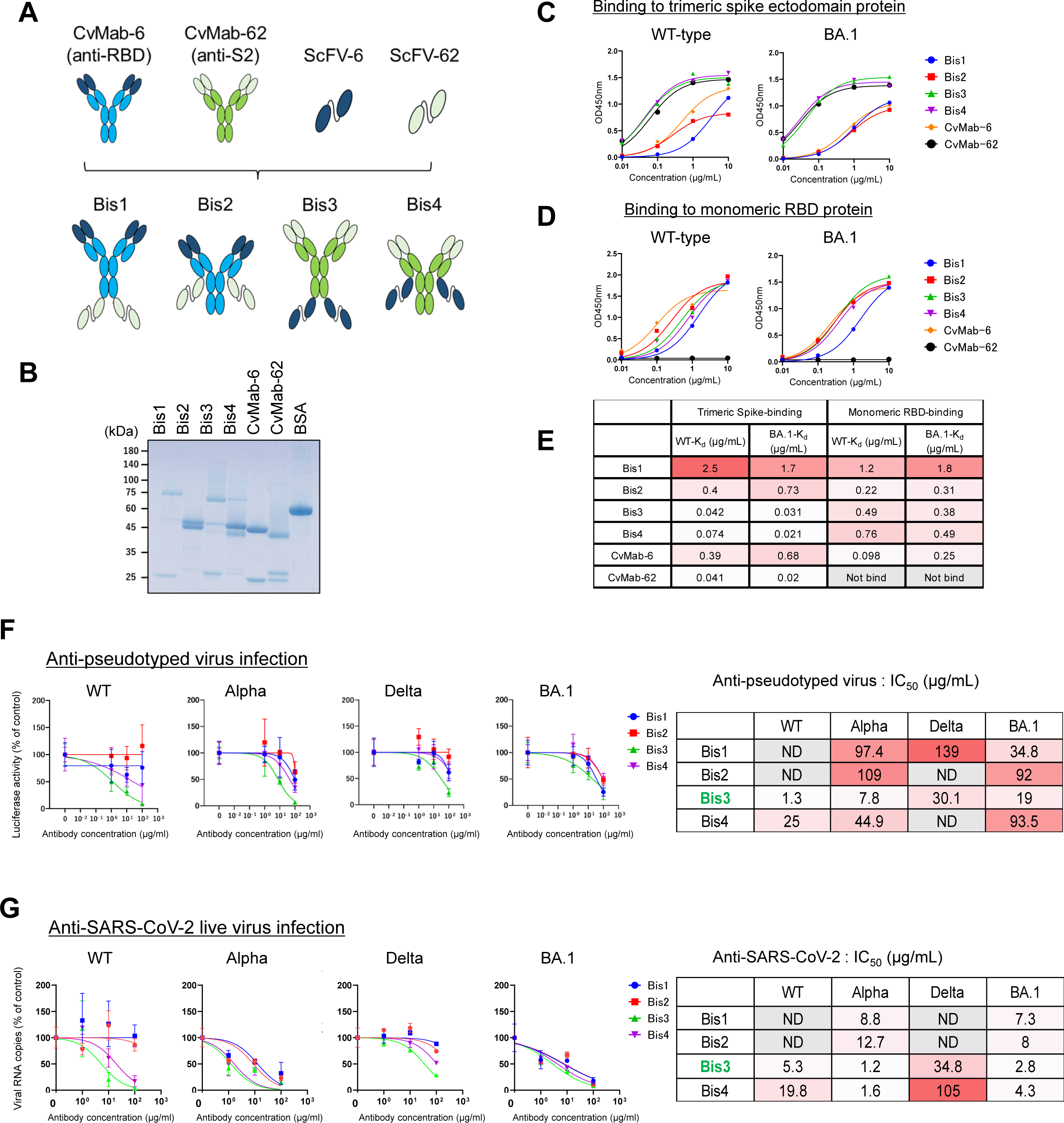
Anti-SARS-CoV-2 effects of bispecific antibodies. (A) Schematic of bispecific antibodies. The scFv of anti-S2 CvMab-62 was fused to the C-terminus of anti-RBD CvMab-6 heavy (Bis1) or light (Bis2) chains. Conversely, the scFv of anti-RBD CvMab-6 was fused to the C-terminus of the anti-S2 CvMab-62 heavy (Bis3) or light (Bis4) chains. (B) The presence of recombinant bispecific antibodies was confirmed by SDS-PAGE, followed by Coomassie blue staining. (C) *In vitro* binding of bispecific antibodies to the trimeric spike ectodomain consists of 1231 amino acids, as measured by ELISA. The trimeric spike ectodomain protein (WT: Wuhan type, or BA.1) was coated in the wells, and bispecific antibodies at the indicated concentrations were added. (D) *In vitro* binding of bispecific antibodies to monomeric RBD consists of 319–541 amino acids measured by ELISA. The RBD protein (WT: Wuhan type, or BA.1) was coated in the wells, and bispecific antibodies at the indicated concentrations were added. (E) Summary table of ELISA. KD values are estimated by GraphPad Prism9. (F) Neutralization of bispecific antibodies against SARS-CoV-2 pseudotyped viruses. Pseudotyped viruses were preincubated with antibodies at the indicated concentrations and then used to infectVeroE6/TMPRSS2 cells. Three days post-infection, cellular luciferase activity was measured to estimate the pseudotyped virus infection ratio (n=3). The inhibitory effects of the bispecific antibodies are shown as IC_50_ values summarized in the table on the right side. ND: not determined. (G) Neutralization activity of bispecific antibodies against authentic SARS-CoV-2 viruses. Authentic SARS-CoV-2 variant viruses were preincubated with antibodies at the indicated concentrations and then used to infectVeroE6/TMPRSS2 cells. At 24 h post-infection, viral genomic RNA in cells was measured by quantitative RT-PCR, and viral replication was shown as the ratio of the control (n=4). The inhibitory effects of the bispecific antibodies are shown as IC_50_ values summarized in the table on the right side. ND: not determined.

### The bispecific antibody Bis3 can inhibit endocytosis-associated viral infection and spike-mediated cell-cell fusion

Regarding the viral entry mechanism, SARS-CoV-2 has two cell entry routes^46^: transmembrane serine protease 2 (TMPRSS2)-dependent cell surface-membrane fusion and TMPRSS2-independent endocytosis (Figure S4). SARS-CoV-2 Omicron variants are thought to be endocytosis pathway dominant types.^47–50^ HEK293/ACE2 cells, expressing little TMPRSS2, possess mainly TMPRSS2-independent endocytic pathways, whereas VeroE6/TMPRSS2 cells^51^ appear to possess both pathways. When the effect of bispecific antibodies on the endocytosis-mediated viral entry pathway was examined in HEK293/ACE2 cells, Bis3 showed an antiviral effect against BA.1 and BA.5.2 pseudotyped viruses (Figure 4A, lower graphs). In contrast, BA.2.75 pseudotyped virus infection in either VeroE6/TMPRSS2 or HEK293/ACE2 was not inhibited by Bis3, suggesting that BA.2.75 spike-mediated infection is resistant to Bis3. Overall, our results indicate that Bis3 inhibits the TMPRSS2-independent endocytic pathway.

**Figure 4.**
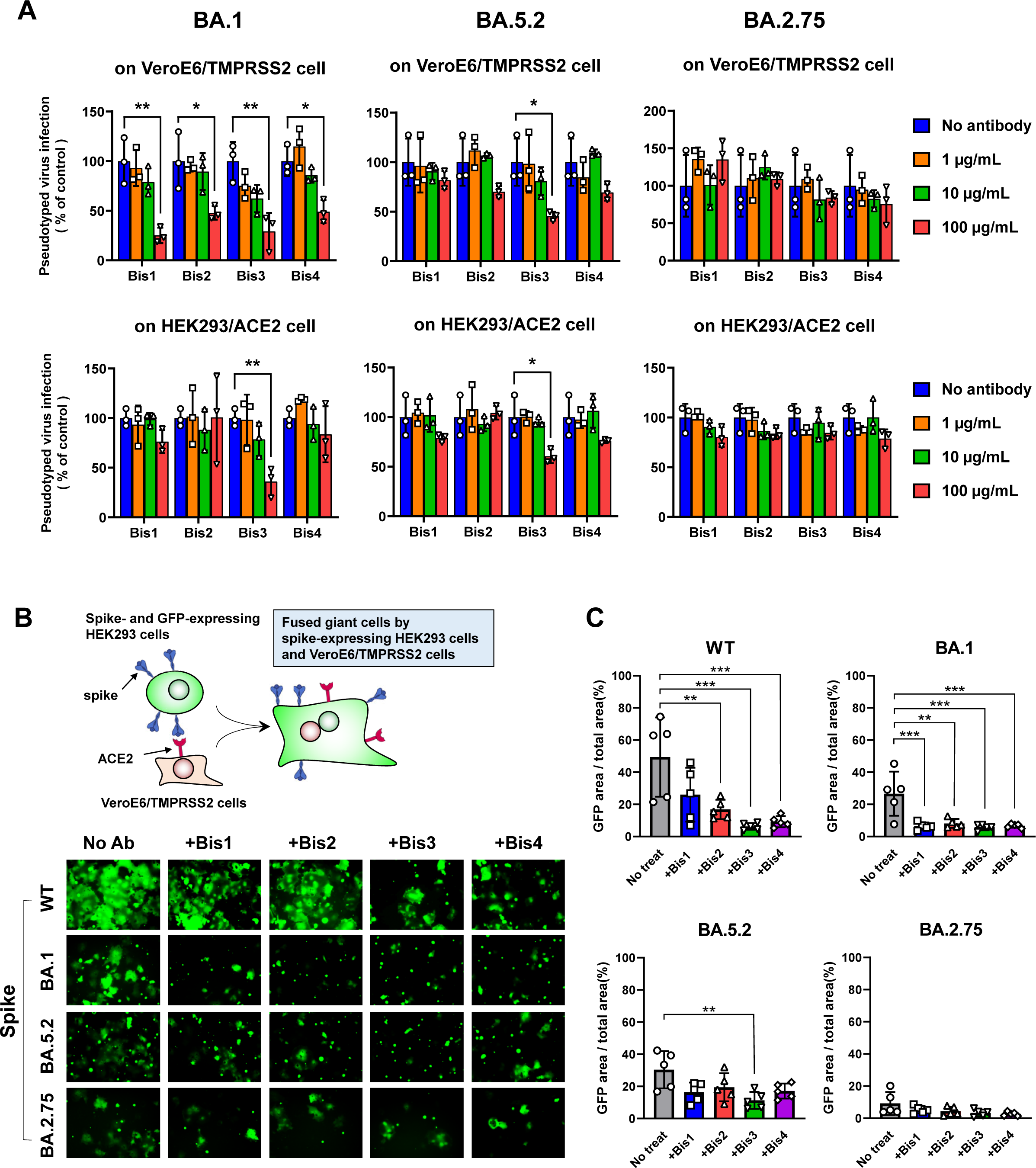
Bis3 suppresses endocytosis-mediated pseudotyped virus infection and spike-mediated cell-cell fusion. (A) Omicron-type SARS-CoV-2 pseudotyped viruses were preincubated with bispecific antibodies at the indicated concentrations and then used to infect VeroE6/TMPRSS2 or HEK293/ACE2 cells. The infection rate was monitored by measuring the luciferase activity of the pseudovirus reporter (n=3). Statistical significance was set at a *p*-value < 0.05, and one-way ANOVA was employed, \**p* < 0.05, \*\**p* < 0.01. (B) Schematics of the cell-cell fusion assay are presented in the upper section. Spike- and GFP-transfected HEK293 cells were suspended, preincubated with bispecific antibodies, and overlaid onto a monolayer of VeroE6/TMPRSS2 cells. After 3h, the GFP-positive fused cells were photographed (lower panels). (C) The green signal area of GFP-positive fused cells, as in B, was quantified using ImageJ, and the results are shown as a bar graph (n=5). Statistical significance was considered at a *p*-value < 0.05, and one-way ANOVA was employed; \*\**p* < 0.01, \*\*\**p* < 0.001.

When two antibodies, an anti-S2 and an anti-RBD antibody, which cannot directly inhibit the binding between the RBD of the spike protein and its receptor ACE2, were combined as Bis3, the inhibitory effect on infection was enhanced. However, the mechanism underlying this enhancement remains unclear. Previous studies have suggested that inhibition of the membrane fusion step mediated by the S2 region of the spike protein is a mechanism for infection inhibition by anti-S2 antibodies.^34^ Therefore, a cell-cell fusion assay was conducted using HEK293 cells expressing the Wuhan-type, Omicron BA.1, BA.5.2, or BA. 2.75 strain spike protein, and VeroE6/TMPRSS2 cells, to investigate the effect of bispecific antibodies on cell membrane fusion. The results demonstrated that each of the four bispecific antibodies showed varying degrees of inhibitory effects on cell-cell fusion. In particular, Bis3 exhibited significantly strong inhibitory activity to cell-cell fusion of Wuhan-type, BA.1, and BA.5.2 (Figures 4B and 4C). Cell-cell fusion induced by the BA.1 spike protein was susceptible to all four bispecific antibodies, consistent with the results of infection inhibition experiments using BA.1 pseudotyped virus and live virus (Figure 3F and 3G). The spike proteins of BA.5.2 showed weak but Bis3-sensitive cell-cell fusion activity, whereas BA.2.75 showed little cell-cell fusion activity in this setting (Figure 4C). The low fusogenic activity observed for BA.2.75 spike protein may potentially exhibit a correlation with resistance to Bis3. Taken together, these results indicate that Bis3 targets the fusogenic activity of spike proteins.

### Bispecific antibody Bis3 does not inhibit TMPRSS2-mediated spike cleavage

SARS-CoV-2 spike-mediated infection involves TMPRSS2-dependent spike protein cleavage into the S1 and S2 portions. We examined the effect of Bis3 treatment on TMPRSS2-dependent spike protein cleavage during cell-cell fusion. In this study, the split green fluorescent protein (GFP) technique was introduced into the cell-cell fusion assay.^52, 53^ When GFP 1-10- and spike-expressing HEK293 cells were fused with GFP11-, ACE2-, and TMPRSS2-expressing HEK293 cells, the GFP signal was restored in the fused giant cells (Figure 5A). The inhibitory effect of Bis3-pretreatment on this GFP-signal ratio was measured, and the results showed that Bis3 significantly inhibited TMPRSS2-dependent and -independent GFP signaling (Figures 5B and 5C). Spike protein cleavage was then examined by western blot analysis, probing the S2 fragment cleaved at the Ser-686 position, showing that Bis3 treatment did not suppress TMPRSS2-dependent S2 fragment production (Figure 5D, upper panels). Collectively, these data indicated that Bis3 did not interfere with TMPRSS2-dependent spike cleavage during cell-cell fusion, suggesting that Bis3 targets the spike S2 fragment-mediated fusion process downstream of TMPRSS2-dependent spike protein cleavage (Figure 5E).

**Figure 5.**
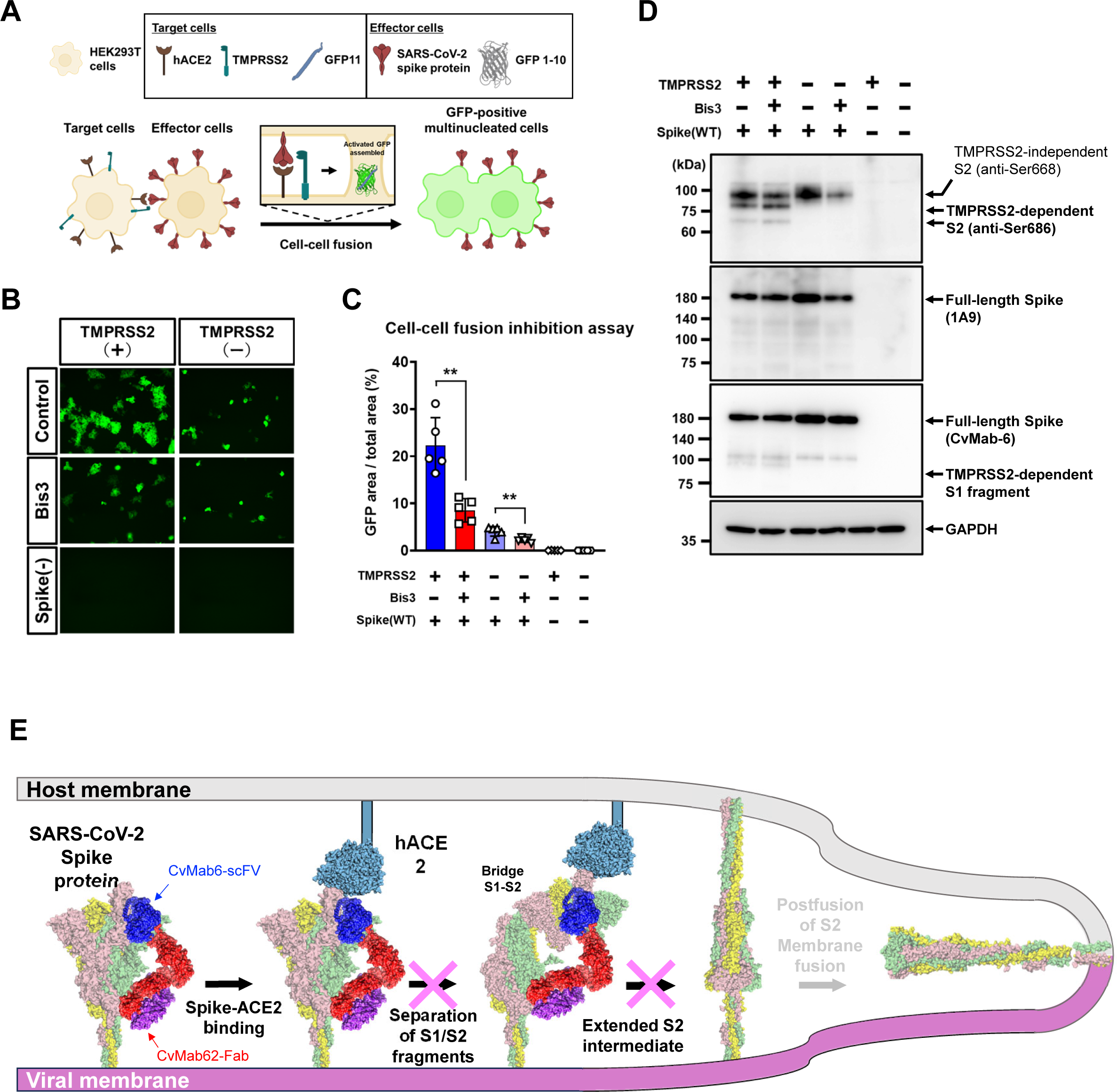
Inhibition of cell-cell fusion by bispecific antibody is independent of S2 cleavage by TMPRSS2. (A) Schematics of the cell-cell fusion assay with a split GFP system are presented. As target cells, HEK293T cells were transfected with ACE2, TMPRSS2 and split GFP11. As effector cells, SARS-CoV-2 spikes and split GFP 1-10 were co-transfected into HEK293T cells. When both cell types were mixed to generate fused cells, reconstituted GFP signals were detected. (B) Wuhan-type spike-expressing effector cells were suspended, preincubated with bispecific antibodies, and overlaid onto a monolayer of target cells. After 3 h, the GFP-positive fused cells were photographed (left panels). (C) The green signal area of GFP-positive fused cells, as in B, was quantified using ImageJ, and the results are shown as a bar graph (n=5). Statistical significance was considered at a *p*-value < 0.05, and one-way ANOVA was employed; \*\**p* < 0.01. (D) After cell fusion by target and effector cells, as in B, S2 cleavage in effector cells was examined by western blot analysis probed with an anti-Ser 686 S2 antibody. TMPRSS2-dependent S2 cleavage was detected and was not affected by bispecific antibody preincubation. (E) Hypothesis of the mechanism of action of the bispecific antibody Bis3. The bispecific antibody Bis3 binds to both the RBD and S2 domains of the spike protein without inhibiting S1/S2 cleavage. Consequently, Bis3 may interfere with the intermediate steps that occur between detachment of the S1 segment and the subsequent membrane fusion process involving the postfusion form of the S2 component.

### Bispecific antibody Bis-Beb restores binding ability to BQ.1.1

Bebtelovimab is a broadly reactive neutralizing antibody effective against many SARS-CoV-2 variants.^54^ However, its efficacy has diminished against recent variants such as BQ.1 and XBB1.5.^27, 30, 31, 55^ Specifically, the effectiveness of bebtelovimab depends on the binding of K444 in the RBD recognition mode (Figure 6A), and it becomes ineffective in cases, such as BQ1.1, with the K444T mutation.^30^ Bispecific antibodies hold promise for overcoming antibody resistance as they can target multiple epitopes. Therefore, to overcome antibody resistance, we developed a novel bispecific antibody, Bis-Beb (Figure 6B). This was accomplished by integrating the antigen-recognition site of bebtelovimab in the form of an scFv into our anti-S2 antibody, CvMab-62, similar to the approach used for Bis3. To validate the binding of this bispecific antibody to the spike protein of BQ1.1, we conducted an ELISA. The Bis-Beb Kd value for the BQ.1.1 trimeric spike ectodomain was more than ten times lower than that of bebtelovimab, whereas the Kd value for the BA4/5 trimeric spike ectodomain was similar to that of the original antibody, bebtelovimab (Figures 6C–6E). Binding of CvMab-62 to the BA4/5 and BQ.1.1 trimeric spike ectodomains were comparable.

**Figure 6.**
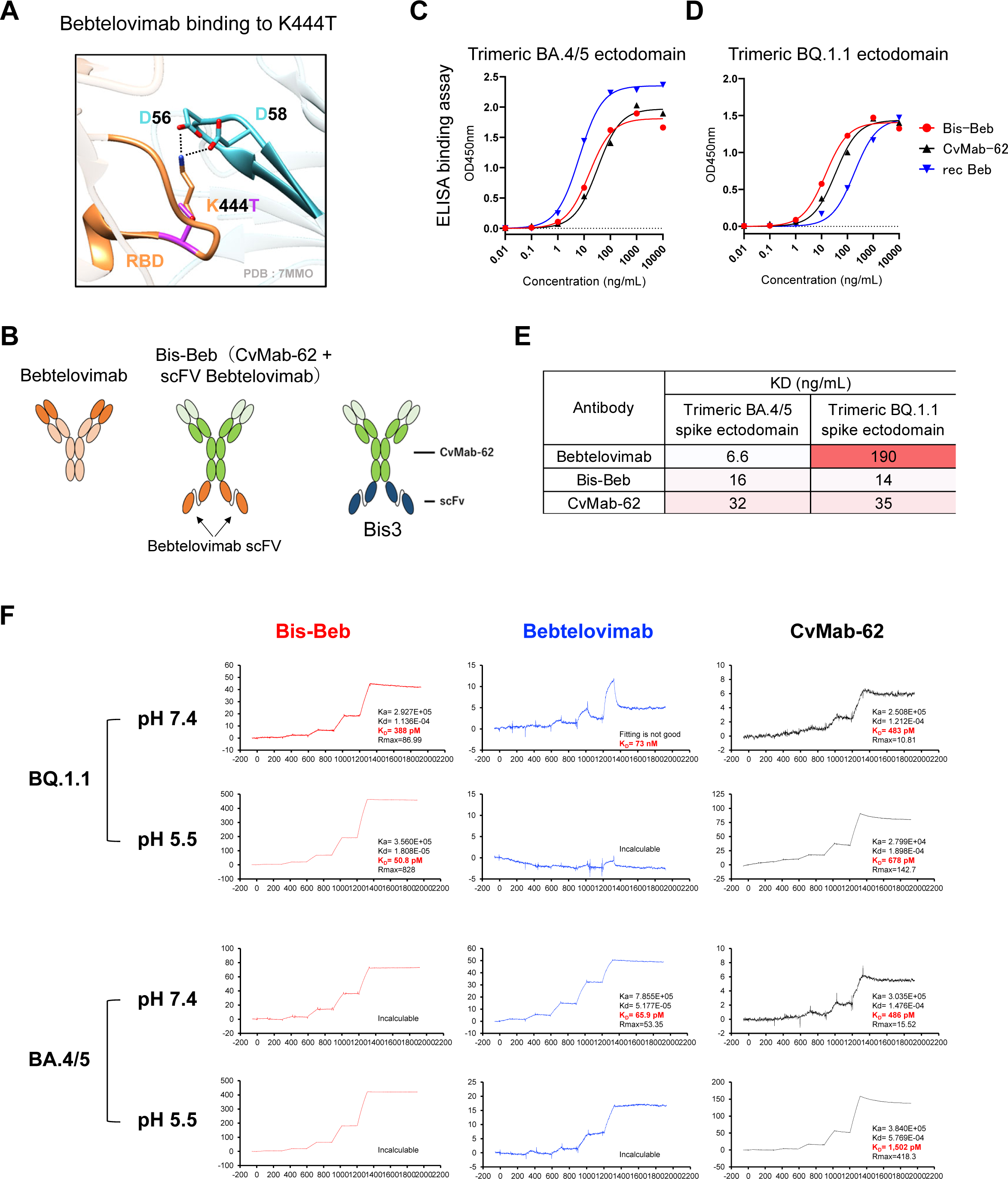
Bispecific antibody constructed with bebtelovimab and CvMab-62. (A) Structural model of bebtelovimab CDR binding to SARS-CoV-2 RBD. The K444 residue in the RBD of the SARS-CoV-2 spike protein interacts with D185 and D187 in the heavy-chain CDR of bebtelovimab. The BQ.1.1 variant of the spike protein has a mutation at K444 (replaced by T), which is responsible for making it resistant to bebtelovimab. (B) Schematic of the bispecific antibody Bis-Beb. The scFv of anti-RBD bebtelovimab was fused with the C-terminus of the anti-S2 CvMab-62 heavy chains. (C) *In vitro* binding of the bispecific antibody to trimeric spike ectodomain of omicron BA.4/5 consists of 1231 amino acids measured by ELISA. (D) *In vitro* binding of the bispecific antibody to the trimeric spike ectodomain of Omicron BQ.1.1, consisting of 1231 amino acids, as measured by ELISA. The wells were coated with the trimeric spike ectodomain protein, and bispecific antibodies at the indicated concentrations were added to evaluate antibody binding to the trimeric spike protein. (E) Summary table of the *in vitro* binding ability of the bispecific antibody in ELISA. The KD values were estimated using C and D by GraphPad Prism9. (F) SPR analysis of the bispecific antibody against the trimeric spike ectodomain. The spike ectodomain, either BQ.1.1 or BA. 4/5 was captured as a ligand, and two buffer conditions, pH 7.4 and pH 5.5, were tested. Antibody as an analyte, Bis-Beb (red lines), bebtelovimab (blue lines), and CvMab-62 (black lines) were tested. The response curves are representative of the two experiments.

Surface Plasmon Resonance (SPR) experiments were conducted to assess the binding strength between Bis-Beb and the spike ectodomain (Figure 6F). In these SPR experiments, we investigated the binding of antibodies with the BQ.1.1 and BA.4/5 trimeric ectodomains under two different pH conditions: neutral (pH 7.4) and acidic (pH 5.5), which served as a representative model for the acidic milieu typically encountered within endosomes. Bis-Beb exhibited strong and stable binding of BQ.1.1 to the BA.4/5 ectodomain under both pH conditions (red lines). In contrast, the original bebtelovimab showed strong binding to the BA4/5 ectodomain, but it rapidly dissociated from the BQ.1.1 ectodomain at pH 7.4. At pH 5.5, it exhibited a complete lack of binding interaction (blue lines). Similar to Bis-Beb, CvMab-62 exhibited strong and stable binding under both conditions (black lines). These results highlight that bebtelovimab has weak binding to BQ.1.1, particularly under acidic conditions, while the bispecific antibody Bis-Beb exhibits strong and stable binding similar to its parent, CvMab-62, towards both BQ.1.1 and BA.4/5.

### Bispecific antibodies overcome bebtelovimab resistance

Figure 6 shows that Bis-Beb exhibited a stronger binding affinity to BQ.1.1, compared with bebtelovimab. ELISA was conducted to investigate whether Bis-Beb inhibits the binding between the BQ.1.1 RBD and ACE2 (Figure 7A). The results showed that both the original bebtelovimab and CvMab-62, even at a concentration of 10 µg/mL, were unable to inhibit the binding between the BQ.1.1 RBD and ACE2. In contrast, Bis-Beb significantly inhibited binding between the BQ.1.1 RBD and ACE2 (Figure 7A, red bar).

**Figure 7.**
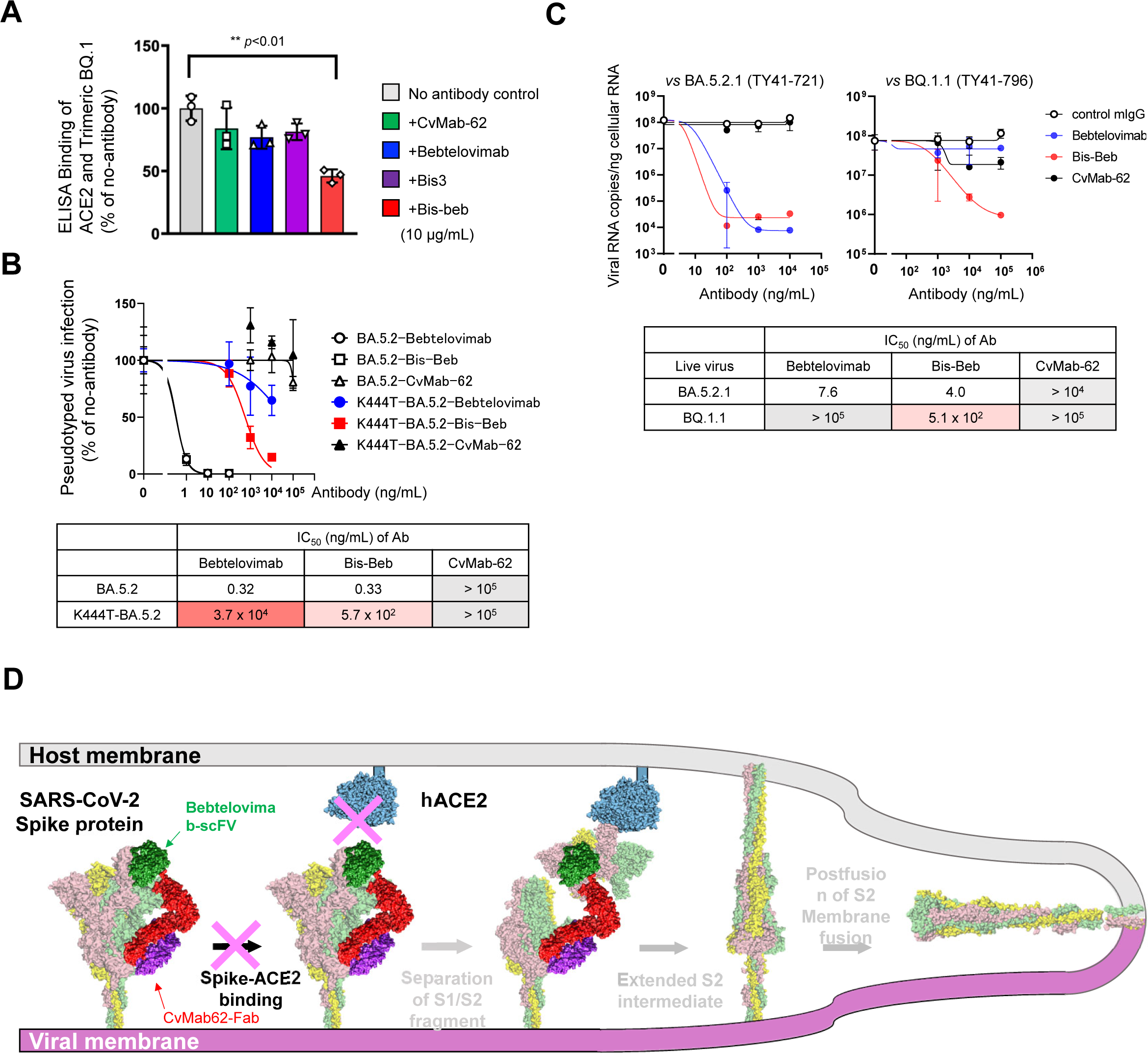
Neutralizing ability of Bis-Beb against bebtelovimab-resistant BQ.1.1. (A) Inhibition of *in vitro* ACE2-spike binding by the bispecific antibody was confirmed using ELISA. His-tagged trimeric spike ectodomain of BQ.1.1 was preincubated with bispecific antibodies (10 µg/mL), and then premixtures added to a well coated with recombinant ACE2 protein. After washing, the ACE2-bound trimeric spike protein was probed with anti-His-tag antibodies. (B) Neutralization of bispecific antibodies against SARS-CoV-2 pseudotyped viruses. Pseudotyped viruses (expressing BA.5.2- or K444T mutated BA.5.2-type spike) were preincubated with antibodies at the indicated concentrations and then used to infect VeroE6/TMPRSS2 cells. Three days post-infection, cellular luciferase activity was measured to estimate the pseudotyped virus infection ratio (n=3). The inhibitory effects of bispecific antibodies are shown as IC_50_ values and summarized in the bottom table. (C) Neutralization of bispecific antibodies against authentic SARS-CoV-2 Authentic SARS-CoV-2 variants (BA.5.2.1 or BQ.1.1) were preincubated with antibodies at the indicated concentrations and then use to infect VeroE6/TMPRSS2 cells. At 24 h post-infection, viral genomic RNA in the cells was measured by quantitative RT-PCR, and viral replication shown as the ratio of the control (n=4). The inhibitory effects of bispecific antibodies are shown as IC_50_ values and summarized in the bottom table. (D) Mechanism of action of the bispecific antibody Bis-Beb. Bis-Beb binds to the RBD and S2 domains of the spike protein. Bis-Beb can restore the ability to inhibit binding between BQ.1.1 RBD and ACE2 and has the capacity to interfere with the subsequent membrane fusion process involving the postfusion form of the S2 component.

Next, we evaluated antiviral activity against BQ.1-type viruses. When testing the BA.5.2 pseudotyped virus, both bebtelovimab and Bis-Beb displayed similar low IC_50_ values (Figure 7B, open circles and open squares, respectively). However, when confronted with the BA.5.2 pseudotyped virus carrying the K444T mutation, a known factor in bebtelovimab resistance,^27, 30, 31^ Bis-Beb exhibited an approximately 60-fold lower IC_50_ value compared to bebtelovimab (Figure 7B; blue circles and red squares, respectively). These findings suggest that Bis-Beb effectively overcomes resistance caused by the K444T mutation.

This inhibitory effect was further confirmed using live viral infection experiments (Figure 7C). When VeroE6/TMPRSS2 cells were infected with BA.5.21 (TY41-721), Bis-Beb demonstrated a low IC_50_ value similar to that of bebtelovimab. Conversely, for BQ.1.1 (TY41-796) infection, bebtelovimab proved ineffective, whereas Bis-Beb exhibited infection-inhibitory activity with submicrogram/mL IC_50_ values (Figure 7C, red symbols). Taken together, these results suggest that Bis-Beb restores the inhibitory effect on the binding between the RBD of BQ.1.1 and ACE2, thereby inhibiting infection with BQ.1.1 (Figure 7D).

## DISCUSSION

In this study, we generated neutralizing bispecific antibodies in the IgG-scFv format by combining ineffective anti-RBD antibodies with anti-S2 antibody, which is a new and distinct structure from previous studies. Notably, the epitope targeted by the anti-S2 antibody in this study was a novel location near the known epitope of anti-S2 antibodies. Furthermore, the structure of the bispecific antibodies, simple fusion of the scFv of the anti-RBD antibody with the C-terminus of the heavy chain of the anti-S2 antibody, can enhance neutralization activity. Moreover, by applying this basic structure, we created a bispecific antibody by combining the scFv of bebtelovimab with our anti-S2 antibody, demonstrating partial overcoming of the resistance to BQ.1.1. This suggests that neutralizing bispecific antibodies, combining existing therapeutic antibodies with S2 antibodies, can revive the value of anti-RBD antibody therapeutics which have diminished in utility owing to resistance issues. Consequently, this approach represents a promising strategy for overcoming antibody therapeutic resistance issues and is worthy of consideration.

Numerous antiviral antibodies with inhibitory activity against SARS-CoV-2 infection have been developed, and some have been applied in clinical settings.^4, 6, 56^ While all anti-SARS-CoV-2 antibody therapeutics target the RBD, newly emerged variants have immune-evasive mutations, particularly within the RBD. Consequently, the clinical application of anti-RBD antibodies has led to reduced neutralization and inhibitory activities.^22, 27, 30, 31, 57^ Although efforts to develop broadly neutralizing antibodies that target pan-coronavirus conserved epitopes are actively underway,^58–65^ the possibility of emerging immune-evading mutant variants remains, posing a recurrent challenge to monoclonal antibody therapy. Therefore, the exploration of alternative approaches is attractive. Another approach is to investigate different antibody formats, bispecific or multispecific, that can simultaneously and synergistically bind to multiple epitopes.^42–44, 66–73^

A study on bispecific antibodies combining anti-RBD and anti-S2 antibodies has been reported.^44^ The structure of these bispecific antibodies involved a combination of scFvs from neutralizing antibodies against RBD and S2, arranged in tandem in the scFv-scFv-Fc format. Unfortunately, the specific epitope targeted by the anti-S2 antibodies has not yet been described. These bispecific antibodies did not show significant improvement in blocking the binding between the RBD and ACE2 compared with the monoclonal antibodies from which they originated. However, their infection-inhibitory activity against mutant variants was enhanced by bispecific antibody formation. This suggests that developing bispecific antibodies targeting both the RBD and S2 is an effective approach for creating broad-spectrum neutralizing antibodies against mutant variants.

Unlike anti-RBD neutralizing antibodies, the infection-inhibitory mechanism of anti-S2 antibodies does not involve the inhibition of binding between ACE2 and RBD; thus, another neutralizing mechanism is involved. The S2 region of the spike protein undergoes significant structural changes between prefusion and postfusion states.^74, 75^ A similar phenomenon in the S2 region has been proposed for the MERS-CoV spike protein.^76, 77^ During the transition from the prefusion to the postfusion form, there is a substantial structural alteration in the linker region between subdomain (SD)3 and hepta-repeat (HR)2, and it is suggested that the correct refolding of this linker region is crucial for forming the central HR1–HR2 six-helix bundle. This bundle brings the viral membrane closer to the host membrane during the late stages of fusion transition.^76^ Dynamic structural changes in the S2 region of SARS-CoV-2 have been observed during membrane fusion and intermediate structures have been elucidated.^78^ Specifically, a model was proposed in which the S2 region was extended, sandwiching the SD3 region between HR1 and HR2, allowing it to fold back and form a postfusion six-helical bundle. Notably, many anti-S2 antibodies targeting SARS-CoV-2 have been reported to exhibit infection inhibitory activity, with several antibodies binding upstream of HR2, the S2 stem helix region within residues 1141–1160 of the spike protein.^34, 36–38, 40, 41, 45, 79^ The alpha helix within residues 1148–1156 (FKEELDKYF) upstream of HR2 is a common epitope for most anti-S2 neutralizing antibodies, and residues F1148, E1151, L1152, D1153, Y1151, and F1156 are directly recognized by the anti-S2 antibody S2P6 complementarity-determining regions (CDRs).^34, 45^ In particular, as suggested by the S2P6 model,^34^ anti-S2 antibodies inhibit the structural conversion of the S2 region necessary for cellular membrane fusion.

Regarding the anti-S2 antibody CvMab-62 epitope, Figure 2 shows that CvMab-62 binding to the S2 region does not require residues 1149–1162 (KEELDKYFKNHTSP); however, the D1146E mutation abolished CvMab-62 binding to S2. Hence, it was inferred that the binding mechanism of CvMab-62 differs from that of typical S2 antibodies, primarily because it does not depend on the epitopes commonly required by anti-S2 neutralizing antibodies, and the orientation of residue D1146, which is essential for CvMab-62 binding, is on the opposite side to where S2P6 binds (Figure S5).^34, 45^ In addition, the bispecific antibody Bis3 did not significantly interfere with the TMPRSS2-dependent spike protein cleavage into the S2 fragment. Therefore, it is presumed that CvMab-62 inhibits the fusion process between the virus and the cell membrane after proteolytic cleavage of the spike protein. Interestingly, the anti-SARS-CoV-2 neutralizing antibody SP1-77 inhibits S1 fragment dissociation from the pre-cleaved S1/S2 complex, thereby blocking the activation of the fusion peptide and membrane fusion.^80^ The bispecific antibody Bis3 might potentially obstruct the structural changes in S2 as S1 separates or create steric hindrance against S1 fragment dissociation from the pre-cleaved S1/S2 complex due to dual binding between the RBD and S2 regions (Figures 5E and 7D).

Bis3 was effective in neutralizing the Omicron variants BA.1 and BA.5.2, particularly by blocking infection through the endosomal pathway. Although CvMab-62 and CvMab-6 appeared to interact equally with the spike proteins of BA.1, BA.5.2, and BA2.75, Bis3 was not effective against BA.2.75, although it is believed to use a similar infection pathway. Previous studies have suggested that the RBD of BA.2.75 binds more strongly to its receptor ACE2 than BA.5.^81^ Additionally, the BA.2.75 spike protein exhibits decreased thermostability and a higher frequency of the RBD being in the “up” conformation under acidic conditions, which suggests enhanced cell entry at low pH through the endosomal pathway.^82^ These robust infectious characteristics may explain why Bis3 was unable to inhibit infection with BA.2.75. Another possibility is that the Bis3-inhibiting mechanism is a fusogenic process involving the spike protein. Previous studies and our data indicate that the Omicron spike protein has low fusogenic activity.^46, 49, 82^ The fusogenic activity of BA.2.75 may play only a small role in its infection process; hence, BA.2.75 spike-mediated infection may be unaffected by Bis3. Recent studies have suggested that SARS-CoV-2 cell entry is instigated by clathrin-mediated endocytosis or Rac1-, Cdc42-, and Pak1-mediated macropinocytosis.^82, 83^ SARS-CoV-2 entry and viral spike-mediated cell-cell fusion mechanisms may rely on different signaling pathways for initiation, and the BA.2.75-spike mediated viral entry mechanism may differ from the Bis3-sensitive cell-cell fusion mechanism. Further investigation is required to confirm these observations.

Bebtelovimab is classified as a class 3 anti-RBD antibody.^82^ Notably, the epitope targeted by bebtelovimab is conserved among many SARS-CoV-2 variants. It has shown efficacy against various mutant variants; however, it has lost its effectiveness against recent variants, such as BQ.1.1 and XBB.1.5.^27, 28, 30^ Specifically, a significant contributor to bebtelovimab resistance in BQ.1.1 is the K444T mutation.^29^ It is presumed that this amino acid mutation interferes with the binding between bebtelovimab and RBD. Further, SPR analysis revealed that bebtelovimab exhibits weak binding and rapid dissociation from BQ.1.1, particularly under acidic conditions such as those found in the endosome, where bebtelovimab fails to bind to the BQ.1.1 spike protein (Figure 6F). Bis-Beb, similar to its parent antibody CvMab-62, maintained stable binding to BQ.1.1, even under acidic conditions. Although bebtelovimab alone cannot inhibit the binding between the RBD of BQ.1.1 and ACE2, its ability to block this interaction is crucially restored when incorporated into the bispecific antibody. Our observations suggested that the scFv derived from bebtelovimab is more likely to be in close proximity to the RBD in a bispecific format. This, in turn, may block the binding between the RBD and ACE2.

Taken together, the bispecific antibody, Bis-Beb, generated in our study, exhibited a remarkable capability to restore neutralization activity against the bebtelovimab-resistant variant BQ.1.1. It should be noted that Bis-Beb in our study apparently overcame the resistance conferred by the K444T mutation in BQ.1.1. However, Bis-Beb is not a universal anti-resistance bispecific antibody. The primary reason for bebtelovimab resistance in XBB.1.5 is the V445P mutation, and we did not observe significant overcoming of V445P-mediated resistance by Bis-Beb (unpublished data). Hence, to overcome bebtelovimab resistance in XBB.1.5, alternative strategies are required, and it is crucial to explore the various structural configurations of bispecific antibodies. This will be essential for further consideration in the development of more effective broad-spectrum bispecific neutralizing antibodies.

## LIMITATIONS OF THE STUDY

This study showed that bispecific antibodies constructed using non-neutralizing anti-RBD and anti-S2 antibodies with different epitopes can gain neutralizing activity against antibody-resistant SARS-CoV-2 variants. We analyzed the biochemical characteristics of bispecific antibodies using an *in vitro* assay. However, the bispecific antibodies described here have limited potency in that the anti-S2 CvMab-62 antibody is a mouse IgG_1_ antibody and is not able to be used for human therapy. Secondly, the structural characteristics of CvMab-62 binding to its epitope near the S2 stem helix remain unknown. A three-dimensional structural analysis is required to clarify the molecular mechanism of CvMab-62 binding to the S2 epitope. Finally, animal model experiments to evaluate the *in vivo* safety and efficacy of these novel bispecific antibodies have not been performed. Thus, humanized bispecific antibodies should be prepared and analyzed to determine their pharmaceutical mechanisms *in vitro* and *in vivo*.

## AUTHOR CONTRIBUTIONS

Conceptualization, Y.K. and K.N.; investigation, T.I., Y.Y., K.S., Y.N., Y.S., M.O., M.K., and K.N.; resources, M.F., T.O., Y.T., and T.W.; writing-original draft preparation, T.I., Y.N., Y.S., M.F., Y.K., and K.N.; writing-review and editing, M. F., Y.K., and K.N.; funding acquisition, Y.Y., M.F., Y.K., and K.N. All authors discussed the results and commented on the manuscript.

## ACKNOWLEDGMENTS

This research was supported in part by an on-campus grant in TUS, funded by donations from “Account for Donations to Develop Vaccine and Medicine to Treat COVID-19” established by the Sumitomo Mitsui Trust Bank (K.N), JSPS KAKENHI JP22K15284 (Y.Y), and by Japan Agency for Medical Research and Development (AMED) under Grant Numbers: 21fk0108568h0001 (Y.Y., K.N), 21fk0108568j0101 (M.F), 21wm0325032s0101 (K.N), 21wm0325032j0201 (M.F), JP23ama121008 (Y.K.), JP23am0401013 (Y.K.), 23bm1123027h0001 (Y.K.), JP23ck0106730 (Y.K.), and JP21am0101078 (Y.K.). We thanks Editage (www.editage.jp) for English language editing.

## DECLARATION OF INTERESTS

The authors declare no competing interests related to the present manuscript.

## STAR*METHODS

### Resource availability

#### Lead contact

Further information or requests should be directed to and will be fulfilled by the lead contact, Kohji Noguchi

#### Materials availability

All materials, except for authentic viruses, in this study can be available upon reasonable request, or through commercially available sources.

#### Data and Code availability

All data are contained within the manuscript.

This paper does not report the original code.

The sources of the datasets supporting the current study are presented in the “key resources table” and “STAR Methods” sections.

Any additional information required to reanalyze the data reported in this paper is available from the lead contact upon reasonable request.

#### Experimental Model and Study Participant details

All animal studies were performed with the approval of The Animal Care and Use Committee of Tohoku University (permit number: 2019NiA-001).

### Method details

#### Cell culture

HEK293T cells (ATCC, CRL-3216) and VeroE6/TMPRSS2 (JCRB, JCRB1819) cells were cultured at 37 ℃ in Dulbecco’s modified Eagle medium (DMEM) supplemented with 7.5% fetal bovine serum (FBS) and kanamycin (50 µg/mL). P3U1 (ATCC, CRL-1597) was cultured in a Roswell Park Memorial Institute (RPMI)-1640 medium (Nacalai Tesque, Kyoto, Japan), with 10% heat-inactivated FBS (Thermo Fisher Scientific, Waltham, MA, USA), 100 units/mL of penicillin, 100 µg/mL of streptomycin, and 0.25 µg/mL of amphotericin B (Nacalai Tesque). HEK293/ACE2 cells were generated by transfection of a human ACE2-DYK-expressing plasmid (Cat# MC_0101086, GenScript Japan, Tokyo, Japan) into HEK293 cells (ATCC, CRL-1573), and stable ACE2-expressing clones were isolated after G418 selection.

#### Spike plasmid construction

The plasmid pUC57-2019-nCoV-S (human), containing synthetic cDNA to express SARS-CoV-2 spike protein with human codon optimization, was purchased from GenScript and cloned into the expression plasmid pcDNA3.1. Mutant spike cDNAs was synthesized using GenScript.^83^

#### SARS-CoV-2 viruses

SARS-CoV-2 viruses, Wuhan strain (2019-hCoV/Japan/TY/WK-521/2020, GISAID ID: EPI_ISL_408667), Alpha variant B.1.1.7 (hCoV-19/Japan/QHN001/2020, GISAID ID: EPI_ISL_804007), Delta variant B.1.617.2 (hCoV-19/Japan/TY11-927/2021, GISAID ID: EPI_ISL_2158617), Omicron variant BA.1 (hCoV-19/Japan/TY38-873/2021, GISAID ID: EPI_ISL_7418017), Omicron variant BA.5.2.1 (hCoV-19/Japan/TY41-704/2022, GISAID ID: EPI_ISL_13241868), and Omicron variant BQ.1.1 (hCoV-19/Japan/TY41-796/2022, GISAID ID: EPI_ISL_15579783) were obtained from National Institute of Infectious Disease (Japan) and handled in biosafety level 3 (BSL-3) facilities.

#### Development of CvMab-6 and CvMab-62

Two 6-week-old female BALB/c mice were purchased from CLEA Japan (Tokyo, Japan) and housed under specific-pathogen-free conditions. All animal experiments were approved by the Animal Care and Use Committee of Tohoku University (permit number: 2019NiA-001). Each BALB/c mouse was intraperitoneally (i.p.) immunized with 100 µg of N-terminal His-tagged S1 spike protein of SARS-CoV-2 (Cat# 230-01102, Ray Biotech, Peachtree Corners, GA, USA) for the development of CvMab-6 or 100 µg of His-tagged S2 spike protein of SARS-CoV-2 (Cat# 230-01103, Ray Biotech) for the development of CvMab-62. Imject Alum (Thermo Fisher Scientific) was used for the first immunization. The procedure included three additional immunization procedures (100 µg/mouse), followed by a final booster i.p. injection (100 µg/mouse) 2 days before harvesting the spleen cells, which were subsequently fused with P3U1 cells using polyethylene glycol 1500 (PEG1500; Roche Diagnostics; Indianapolis, IN, USA). Hybridomas were grown in RPMI medium supplemented with hypoxanthine, aminopterin, and thymidine for selection (Thermo Fisher Scientific). The culture supernatants were screened using ELISA to detect SARS-CoV-2 S1 for the development of CvMab-6 or SARS-CoV-2 S2 for the development of CvMab-62. Clone CvMab-6 or clone CvMab-62 culture supernatants in hybridoma-SFM medium (Thermo Fisher Scientific) were purified using an Ab-Capcher (ProteNova, Kagawa, Japan).

#### Development of bispecific antibody

To generate each bispecific antibody, we first constructed an scFv of CvMab-6 (Cv6-scFv) or CvMab-62 (Cv62-scFv) by connecting the V_H_ and V_L_ cDNA of CvMab-6 or CvMab-62 with a linker sequence (GGGGSGGGGSGGGGS). Each scFv cDNA was further fused at the 3′ end of the heavy or light chain cDNA of CvMab-6 or CvMab-62, to create Bis1 (Cv62-scFv fused to the heavy chain of CvMab-6), Bis2 (Cv62-scFv fused to the light chain of CvMab-6), Bis3 (Cv6-scFv to the heavy chain of CvMab-62), or Bis4 (Cv6-scFv to the light chain of CvMab-62). The cDNA of each heavy and light chain was transduced into ExpiCHO-S cells using the ExpiCHO Expression System (Thermo Fisher Scientific). Each antibody was purified using an Ab-Capcher. For bispecific antibodies, the amino acid sequence can be conditionally disclosed upon request.

#### ELISA for antibody development

His-tagged S1 spike protein of SARS-CoV-2 (100 µg) or 100 µg of His-tagged S2 spike protein of SARS-CoV-2 was immobilized on Nunc Maxisorp 96-well immunoplates ((Cat# 439454, Thermo Fisher Scientific) at 1 µg/mL for 30 minutes at 37 °C. After washing with phosphate buffered saline (PBS) containing 0.05% Tween20 (PBS-T; Nacalai Tesque), wells were blocked with 1% bovine serum albumin (BSA) in PBS-T for 30 minutes at 37 °C. The plates were incubated with primary antibodies followed by peroxidase-conjugated anti-mouse immunoglobulin (1:1000; Agilent Technologies, Santa Clara, CA, USA). Finally, enzymatic reactions were performed using an ELISA POD substrate TMB kit (Cat# 05298-80, Nacalai Tesque). The absorbance at 655 nm was measured using an iMark microplate reader (Bio-Rad Laboratories, Berkeley, CA).

#### CvMab-6 epitope mapping

The 22 peptides from the S1-RBD sequence were synthesized using PEPScreen (Sigma-Aldrich, St. Louis, MO, USA). The cysteine in each peptide was converted into serine. The peptide sequences are listed in Table 1. The binding assay was performed using ELISA, as described above. Briefly, each peptide was immobilized on Nunc Maxisorp 96-well immunoplates at 1 µg/mL for 30 min at 37 °C. After washing with PBS-T, wells were blocked with 1% BSA in PBS-T for 30 min at 37 °C. The plates were then incubated with CvMab-6, followed by incubation with peroxidase-conjugated anti-mouse immunoglobulins (1:1000). Finally, enzymatic reactions were performed using an ELISA POD substrate TMB kit (Nacalai Tesque, Inc.). The absorbance was measured at 655 nm using an iMark microplate reader.

#### CvMab-62 epitope mapping

S2 deletion mutant cDNAs encoding the SARS-CoV-2 spike protein S2 region, as shown in Table 2, were cloned into the pUC57 plasmid (GenScript Japan). Recombinant proteins were expressed by IPTG induction in *E.coli* JM109, and cell lysates were prepared. Anti-S2 antibody reactivity to these S2 proteins was examined by western blot analysis using CvMab-62 and 1A9 (Cat# GTX632604, GeneTex, CA, USA) and a polyclonal anti-S2 antibody (Cat# 40590-T62, Sino Biological, Beijing, China).

#### Pseudotyped virus neutralization assay

Retrovirus-based pseudotyped virus production was performed as previously described.^84^ Briefly, phCMV-Gag-Pol 5349 and reporter pTG-Luc126 plasmids^85^ were co-transfected into HEK293T cells along with SARS-CoV-2 spike expressing plasmids using the PEIpro® transfection reagent (Cat# 101000017, Polyplus Transfection, New York, NY). Medium was added the day after transfection. The cell supernatant containing pseudotyped virus was collected 72 h post-transfection, filtered through a 0.45 µm filter, and aliquoted to be stored at −80 ℃.

VeroE6/TMPRSS2 cells were seeded in 96-well white plates. The next day, antibodies were serially diluted in medium and mixed with pseudotyped viruses for 1 h at 37 ℃, and then added to the wells. After 3 days, the medium was removed. Cells were washed once with PBS and subsequently lysed using a luciferase assay reagent (Cat# MLT100, PicaGene Meliora Star-LT Luminescence Reagent; TOYO B-NET, Tokyo, Japan). Transduction was performed in triplicate in each experiment. The average and standard deviation (SD) were calculated, and reproducibility was confirmed by at least two independent biological experiments.

#### Authentic SARS-CoV-2 neutralization assay

VeroE6/TMPRSS2 cells were cultured in DMEM (Cat# 044-29765, Fujifilm Wako Pure Chemical Industries, Osaka, Japan) supplemented with 10% (v/v) heat-inactivated FBS (Cat# FBS-12A, Capricorn Scientific GmbH, Ebsdorfergrund, Germany), 100 U/ml penicillin G, 100 µg/ml streptomycin sulfate (Cat# 26253-84, Nacalai Tesque), and 1 mg/ml G418 (Cat# 09380-86, Nacalai Tesque). One day before SARS-CoV-2 infection, the cells were seeded in a 48-well plate (Cat# 3548, Corning, Glendale, AZ, USA) at a density of 5 × 10^4^ cells/well. Viruses (WT, 0.001 TCID50/cell; Alpha, 0.01 TCID50/cell; Delta, 0.1 TCID50/cell; Omicron BA1, 0.1 TCID50/cell; and Omicron BA5.2.1, 0.001 TCID50/cell) were preincubated with CvMab-62, Bis1, Bis2, Bis3, Bis4, bebtelovimab, Bis-Beb, or mouse IgG (Cat# 140-09511, Fujifilm Wako Pure Chemical Industries; Cat# 1015-000-003, Jackson ImmunoResearch, West Grove, PA, USA) at 37 ℃ for 30 min. The mixtures were added to the cell monolayers and the cells incubated for 2 h. After removing the antibody-virus mixtures, the cells were washed with PBS and cultured in normal growth medium in the presence of antibodies for 24 h. Viral RNA copies in the culture supernatant and cells were determined as described below.

#### Quantification of viral RNA

Total RNA from the culture supernatant and cells was extracted using the Viral RNA/Viral Nucleic Acid Mini Kit (Cat# FAVNK 001–2, Favorgen Biotech, Pingtung City, Taiwan) and the Tissue Total RNA Purification Mini Kit (Cat# FATPK 001–2, Favorgen Biotech), respectively, according to the manufacturer’s instructions. Viral RNA copies were quantified via real-time RT-PCR analysis using the THUNDERBIRD® Probe One-step qRT-PCR Kit (Cat# QRZ-101, Toyobo, Osaka, Japan). SARS-CoV-2-specific primers (NIID_2019-nCOV_N_F2; 5′-AAATTTTGGGGACCAGGAAC-3′, NIID_2019-nCOV_N_R2; 5′-TGGCAGCTGTGTAGGTCAAC-3′) and probe (NIID_2019-nCOV_N_P2;5′-FAM-ATGTCGCGCATTGGCATGGA-BHQ-3′) were purchased from Eurofins Genomics (Tokyo, Japan).

#### Immunoblotting

The cells were lysed using radioimmunoprecipitation assay (RIPA) buffer (Cat# 16488-34, Nacalai Tesque) containing protease inhibitors. Proteins were separated using SDS-PAGE and transferred to Immobilon-P membranes (Cat# IPVH00010, EMD Millipore, Billerica, MA, USA). After blocking with 5% milk in PBS-T for 1 h, membranes were incubated with primary antibodies at 4 ℃ overnight. Membranes were then washed with PBS-T and incubated with secondary antibodies for 2 h. Membranes were washed again with PBS-T and immunoblot signals were developed using EzWestLumi plus® (Cat# WES-7120, ATTO, Tokyo, Japan) and recorded using an ImageQuant LAS4000 mini-image analyzer (GE Healthcare Japan, Tokyo, Japan). The antibodies used were as follows: anti-spike antibody 1A9 (Cat# GTX632604, GeneTex, CA, USA), anti-cleaved spike (Ser 686) antibody (Cat# 84534, Cell Signaling Technology, Massachusetts, USA), anti-GAPDH 3H12 (Cat# 171-3, Medical & Biological Laboratories, Aichi, Japan), goat anti-mouse IgG antibody conjugated with horseradish peroxidase (Cat# SA00001-1, Proteintech Group, Rosemont, USA), donkey anti-rabbit IgG antibody conjugated with horseradish peroxidase (Cat# NA934, Cytiva, Tokyo, Japan).

#### Immunofluorescence microscopic analysis

HEK 293T cells were seeded in a poly D-lysine-coated four well slide chamber (Cat# 354114, Corning, Corning, NY, USA) and transfected with the SARS-CoV-2 spike protein the following day. Forty-eight hours after transfection, the cells were fixed in 4% paraformaldehyde (Cat# 09154-14, Nacalai tesque, Inc.) for 10 min and blocked with 1% BSA/PBS for 30 min. The cells were then incubated with primary antibodies (CvMab-6 and CvMab-62) at a concentration of 10 µg/mL in 1% BSA/PBS for 2 h at room temperature and washed three times with PBS. Next, the cells were incubated with goat anti-mouse IgG (H+L) cross-adsorbed Alexa Fluor® 488 (Cat# A-11001, Thermo Fisher Scientific, Carlsbad, CA) diluted × 2000 in 1% BSA/PBS at room temperature for 1 h and washed again with PBS three times. Vectashield Vibrance Antifade Mounting Medium (Cat# H-1700, Vector Laboratories, CA, USA) was added, and the cells were incubated at room temperature for 1 h. Images were captured using a BZ-9000 microscope (Keyence, Osaka, Japan). Two-dimensional TIFF images were merged using Adobe Photoshop CS4 Extended software (Adobe Systems Inc., San Jose, CA, USA).

#### Cell-cell fusion assay

To prepare the effector cells, HEK 293T cells were co-transfected with EGFP and SARS-CoV-2 spike protein-expressing plasmids. Cells were collected 48 h after transfection and treated with antibodies at a final concentration of 100 µg/mL at 37 ℃ for 30 min, then added to VeroE6/TMPRSS2 cells as target cells and co-cultured at 37 ℃ for 3 h. After incubation, five fields were randomly selected in each well, and images were captured using a fluorescence microscope BZ-800 (Keyence). Images were analyzed using ImageJ software to quantify the GFP area. To assess TMPRSS2 dependence of cell-cell fusion inhibition, we used HEK293T cells co-transfected with split GFP1-10 and SARS-CoV-2 spike-expressing plasmids as effector cells and HEK293T cells co-transfected with split GFP11 and TMPRSS2 expressing plasmids as target cells, 48 h after transfection.^52^ The remaining steps were performed as described previously. After the cell-cell fusion assay, cell lysates were collected and used for western blotting to confirm the TMPRSS2-dependent cleavage of the spike protein.

#### ELISA for binding affinity calculation

Wells of 96-well microtiter plates were coated with 1 µg/mL purified recombinant SARS-CoV-2 Trimeric Spike ECD and RBD ((Spike WT; Cat# 40589-V08H4, Spike BA.1; Cat# 40589-V08H26, Spike BA.4/5; Cat# 40589-V08H32, Spike BQ.1; Cat# 40589-V08H41, RBD BQ.1; Cat# 40589-V08H143, Sino Biological, Beijing, China) at 4 ℃ overnight. Plates were blocked with 1% BSA in PBS-T for 1 h. Antibodies were diluted in blocking buffer, added to the wells, and incubated for 2 h at room temperature. The bound antibodies were detected using goat anti-mouse IgG antibody conjugated with horseradish peroxidase (Proteintech Group) and TMB substrate (Cat# 34028, 1-Step™ TMB ELISA Substrate Solutions, Thermo Fisher Scientific, Carlsbad, CA). Color development was monitored, 2 M sulfuric acid was added to stop the reaction, and the absorbance was measured at 450 nm using a multi-mode plate reader SpectraMax iD3 (Molecular Devices, CA, USA).

#### ELISA for *in vitro* RBD and ACE neutralization assay

Wells of 96-well microtiter plates were coated with 1 µg/mL recombinant hACE2-Fc^83^ at 4 ℃ overnight. Plates were blocked with 1% BSA in PBS containing 0.05% PBS-T for 2 h. Antibodies and recombinant His-tagged SARS-CoV-2 Spike protein RBD^84^ were mixed and incubated for 1 h at room temperature. The antibodies and RBD mixture were added to the wells and incubated for 2 h at room temperature. RBD bound to ACE2 were detected using Anti-His-tag mAb-HRP-DirecT (Cat# D291-7, Medical & Biological Laboratories, Tokyo, Japan) and TMB substrate (1-Step™ TMB ELISA Substrate Solutions, Thermo Fisher Scientific, Carlsbad, CA). Color development was monitored, 2 M sulfuric acid was added to stop the reaction, and the absorbance was measured at 450 nm using a multi-mode plate reader SpectraMax iD3 (Molecular Devices).

#### The surface plasmon resonance (SPR) experiments

To measure the affinity of each antibody for the antigen (trimeric BQ.1.1 and BA. 4/5 spike ectodomains), we performed SPR analysis using Biacore X100 (Cytiva, Marlborough, MA, USA). HBS-EP + buffer (10 mM HEPES [pH 7.4], 150 mM NaCl, 3 mM EDTA, and 0.05% v/v surfactant P20) and MBS-EP + buffer (10 mM MES [pH 5.5], 150 mM NaCl, 3 mM EDTA, and 0.05% v/v surfactant P20) were used as running buffers. One hundred glycine HCl (pH 1.5) and 50 mM NaOH were used as the dissociation buffer. Anti-His tag mAb (Cat# D291-3, Medical & Biological Laboratories) was immobilized on a Sensor Chip CM5 (Cat# BR100399, Cytiva) using an amine coupling kit ((Cat# BR100050, Cytiva) according to the manufacturer’s standard amine coupling protocol. The level of immobilized trimeric spike ectodomain in active flow cells reached approximately 500 response units. Antibodies were serially diluted (0.11, 0.33, 1.0, 3.0, and 9.0 µg/ml) in the running buffer. The single-binding cycles were injected sequentially with increasing concentrations over both the ligand and reference surfaces. The reference surface, which was an unmodified flow cell, was used to correct systematic noise and instrumental drift. To determine ka, kd, and KD values, the sensorgrams were globally fitted using a 1:1 binding model and analyzed using Biacore X100 Evaluation Software.

#### Structural modeling of the spike protein

Structural modeling was performed using the 3D mol.js software (https://3dmol.csb.pitt.edu/), UCSF Chimera,^86^ and Jalview.^87^ Modeling of ‘SARS-CoV-2 Spike RBD bound with ACE2 (6M0J)’ was performed in 3D mol.js software and used in Figure 2D. Meanwhile, modeling of ‘LY-CoV1404 against SARS-CoV-2 RBD (7MMO)’ and ‘SARS-CoV-2 Spike protein prefusion form (6XR8) and postfusion form (6XRA)’ were performed in UCSF Chimera and used in Figure 8 and S4.

### Quantitative and statistical analysis

Inhibition concentrations (IC_50_ values) during the neutralization assays and KD values in the ELISA binding assays were determined. Data visualization and statistical analyses were performed using GraphPad Prism 9 (GraphPad Software, Inc., Boston, MA, U.S.A). Statistical differences were determined using an unpaired t-test, or a one-way ANOVA followed by the Bonferroni test post hoc test (group ≥3). Data are presented as the means ± SD from triplicated samples, minimum, and reproducibility was confirmed by a minimum of two independent biological experiments. Statistical significance was defined as a *p*-value < 0.05, \**p* < 0.05, \*\**p* < 0.01, and \*\*\**p* < 0.001. Multiple sequence alignments of spike proteins were performed using Clustal Omega software (https://www.ebi.ac.uk/Tools/msa/clustalo/). Quantification of the GFP signal images was performed using ImageJ software (NIH). Where applicable, the statistical parameters are reported in the figure legends.

## SUPPLEMENTAL INFORMATION

**Table S1.**
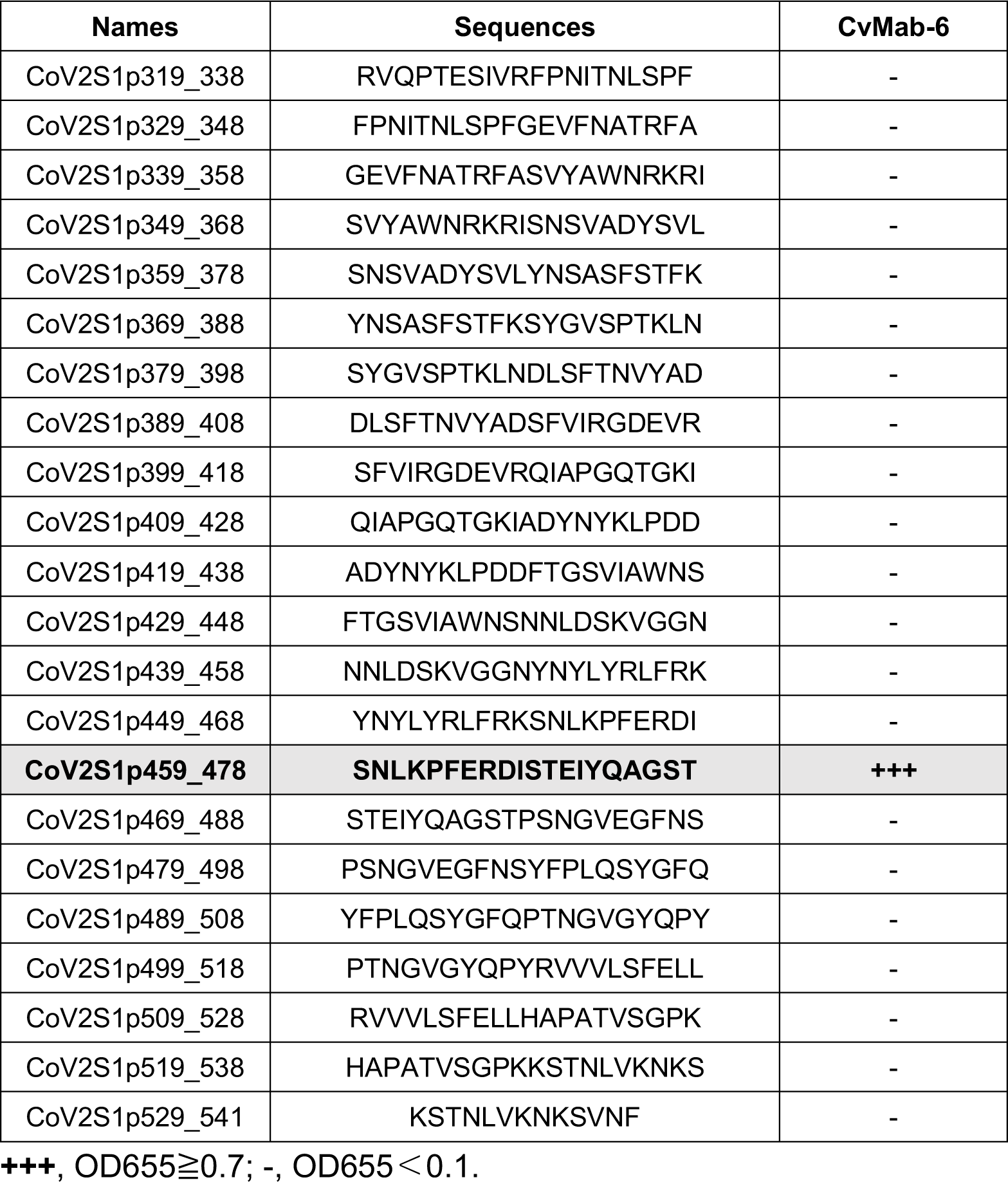
Epitope mapping using PEPScreen.

**Figure S1.**
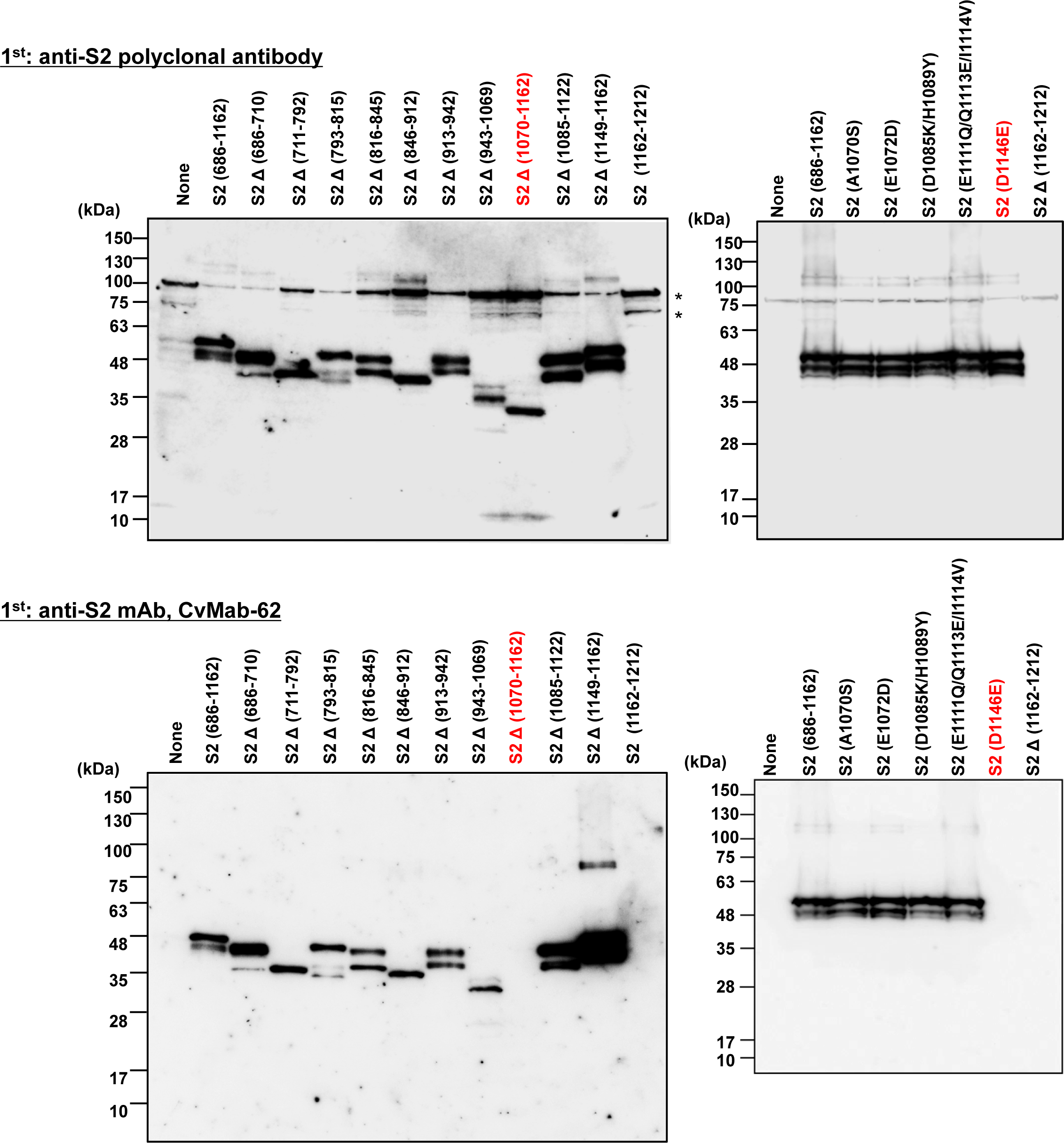
CvMab-62 reactivities to S2 deletion mutants. Western blot analysis using S2 mutants and anti-spike S2 polyclonal antibody and CvMab-62. * indicates non-specific bands.

**Figure S2.**
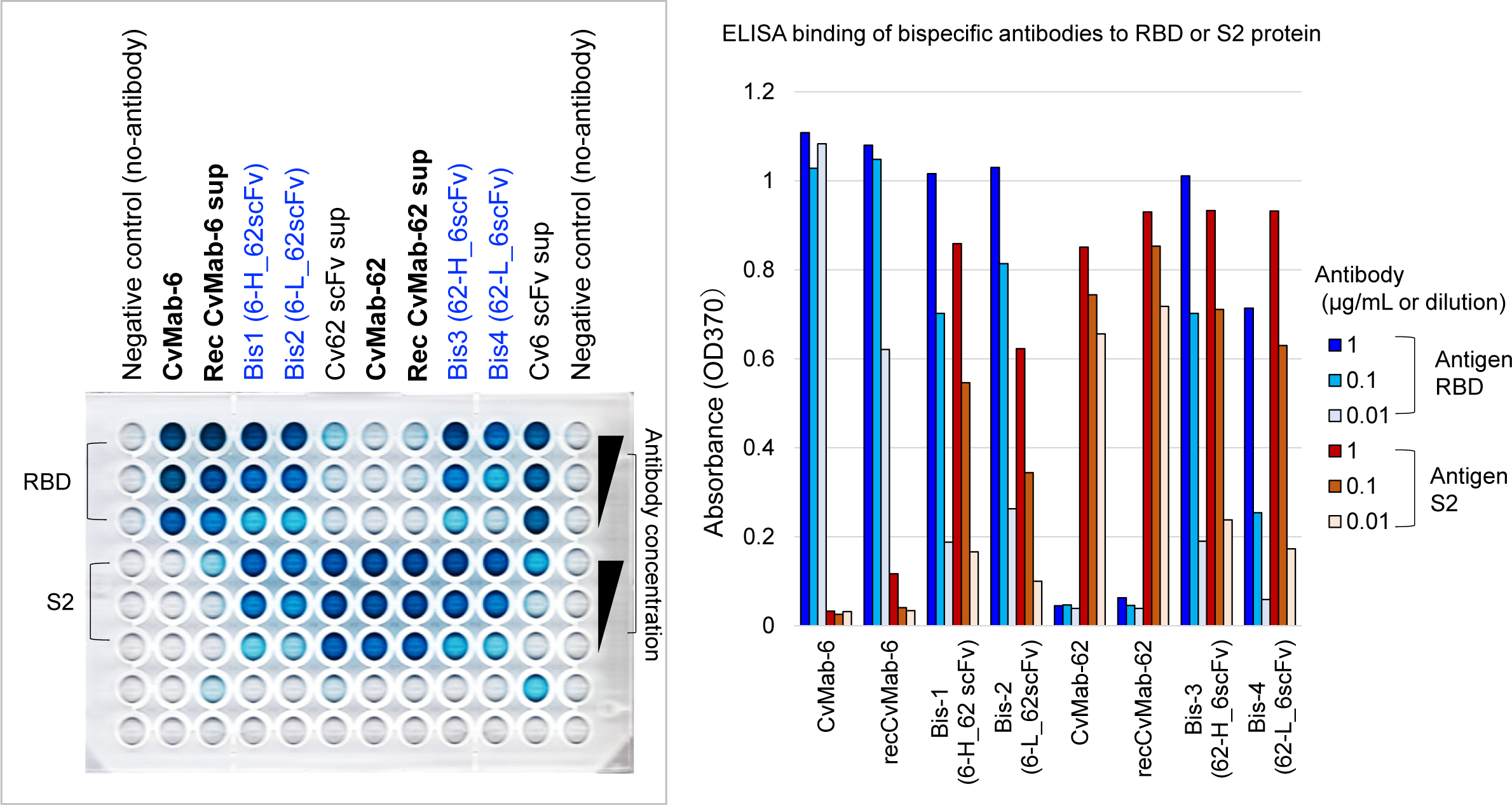
In vitro binding of recombinant bispecific antibodies were confirmed by ELISA. Photographic image of ELISA plate after adding of substrate development is shown at left, and results is summarized as bar graph at light. Binding to RBD and S2 are shown in blue and brawn colored bar graphs, respectively.

**Figure S3.**
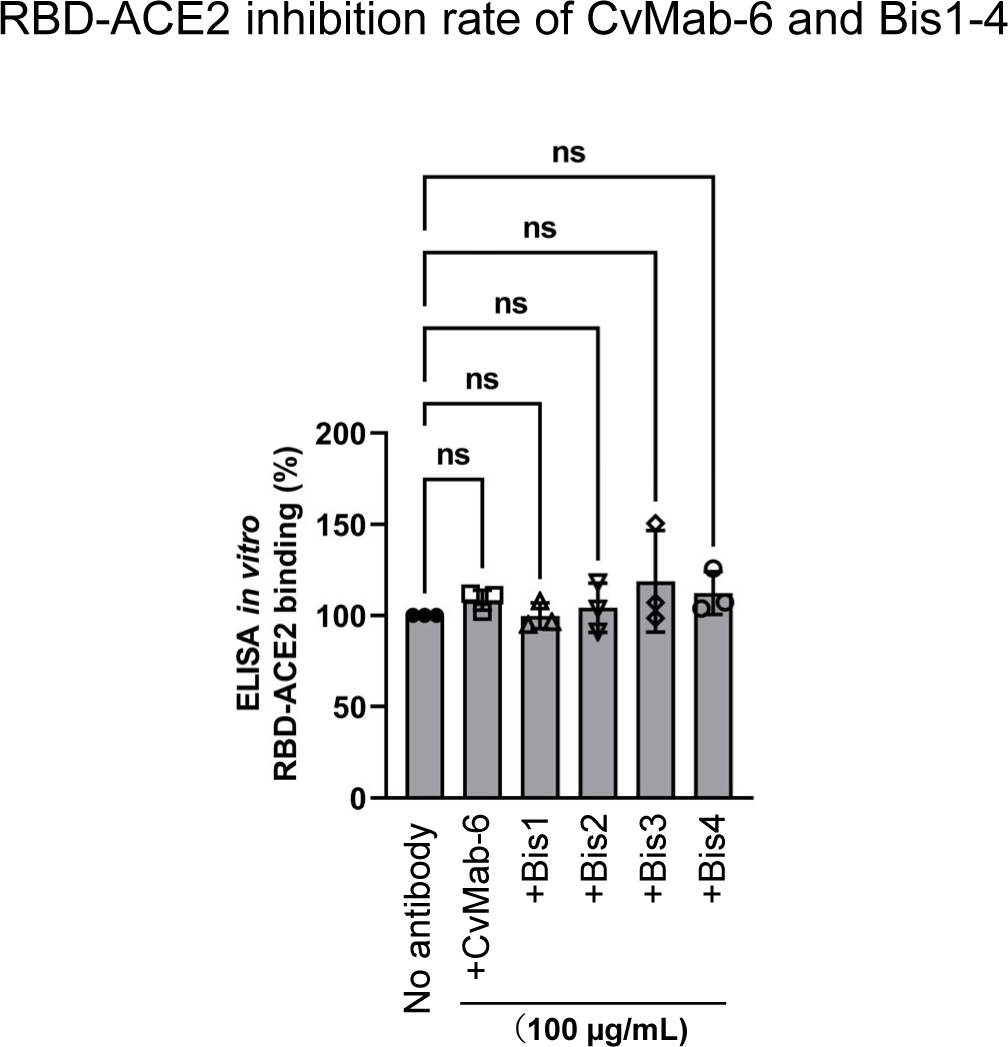
No inhibition of ACE2-RBD binding by CvMab-6 and bispecific antibodies confirmed by ELISA. Recombinant monomeric avi-tag-fused RBD protein was preincubated with bispecific antibodies (100 µg/mL) for 1 h, and then premixtures were added to a well coated with recombinant ACE2 protein for 1 h. After washing well, the ACE2-bound RBD was probed by anti-avi tag antibodies.

**Figure S4.**
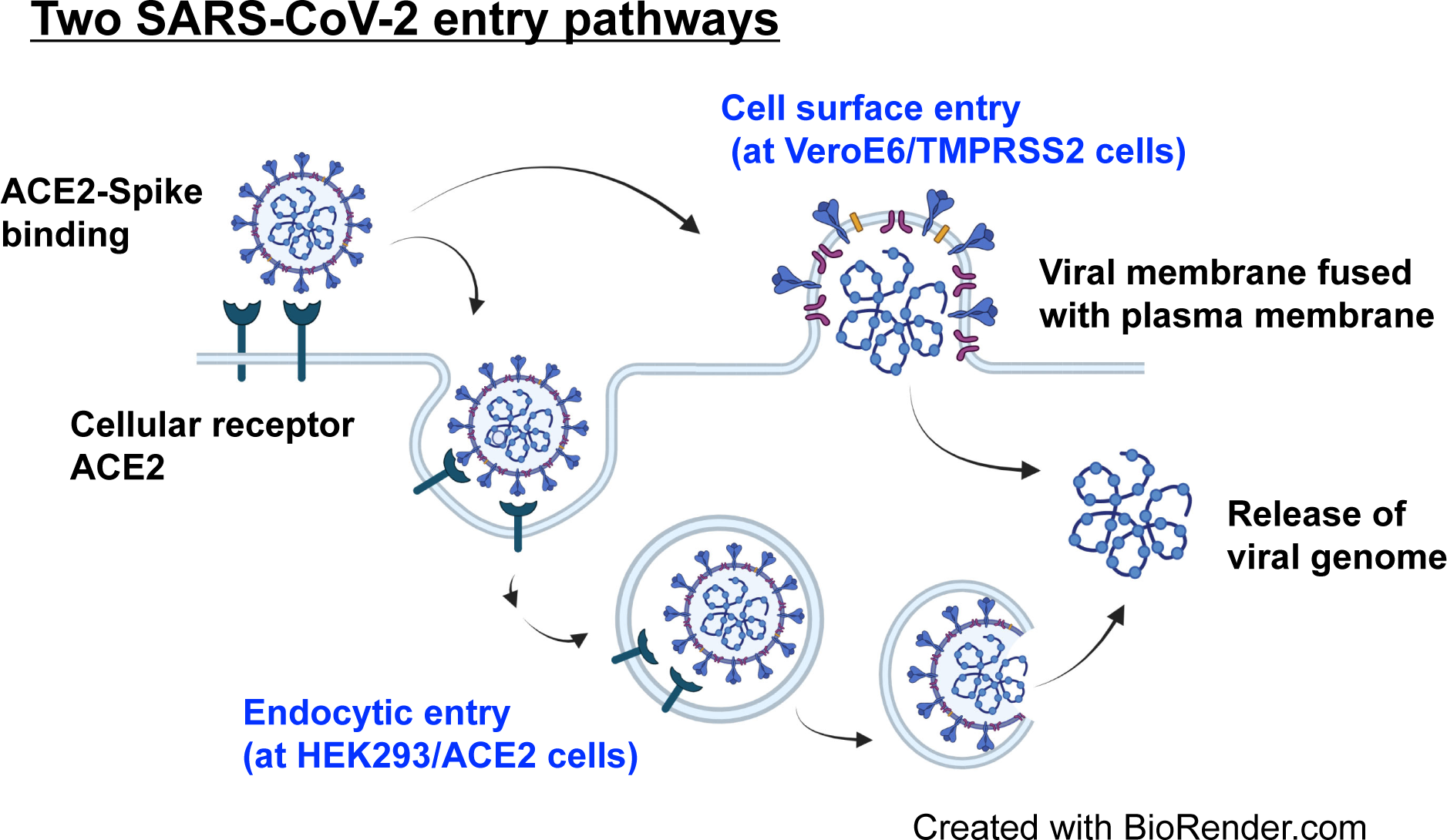
Schematics of SARS-CoV-2 cell entry pathway. SARS-CoV-2 has two cell entry mechanisms, one is cell surface entry via fusion with plasma membrane, and the other is endocytosis-mediated entry. Endocytosis pathway is major SARS-CoV-2 entry route in HEK293/ACE2 cells lacking TMPRSS2, but VeroE6/TMRPSS2 cells has two pathway. This illustration is created with BioRender.com.

**Figure S5.**
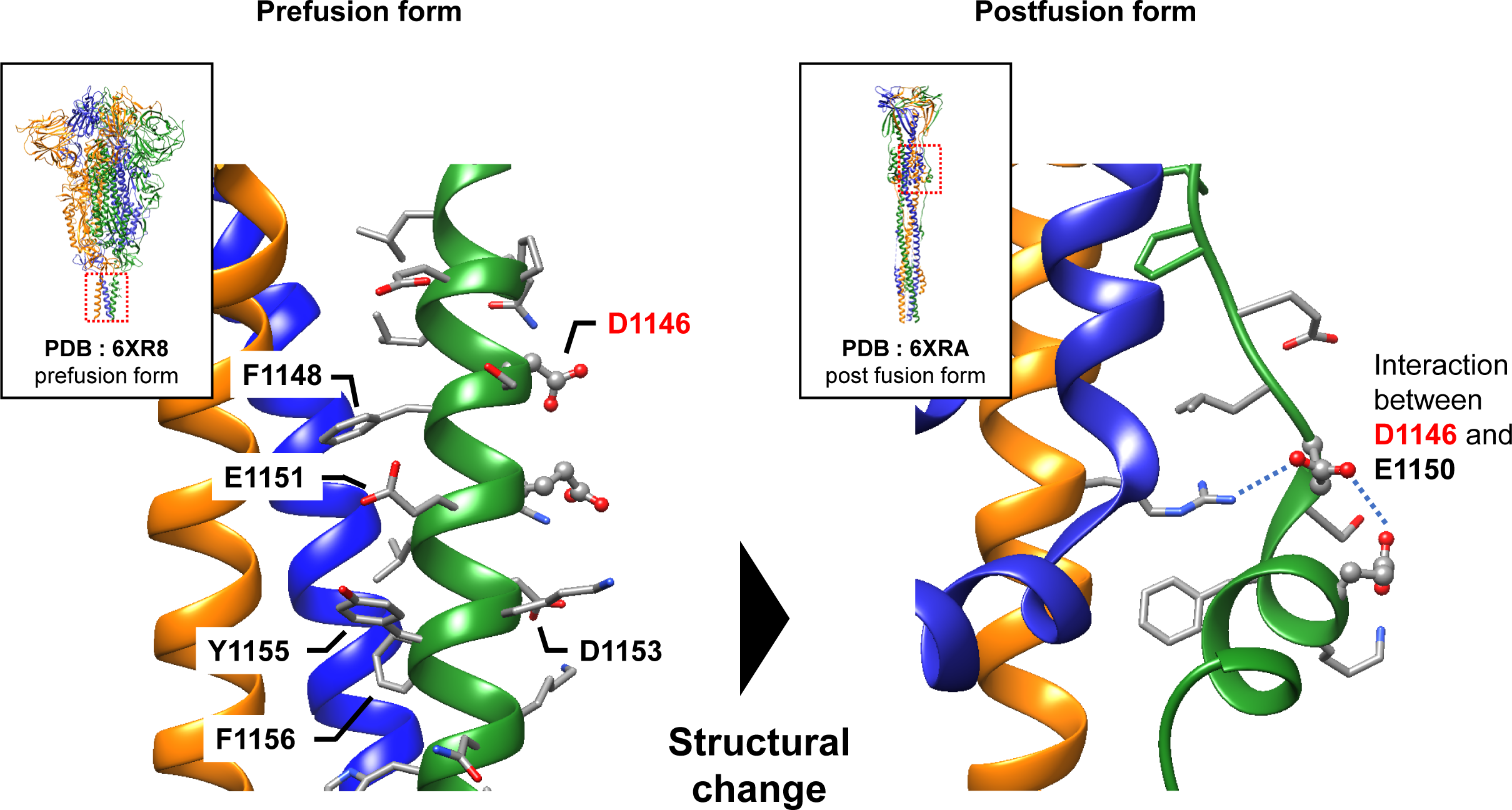
S2 stem helix structural model. The D1146 residue, which impacts the binding of CvMab-62, is located on the opposite side compared to where another anti-S2 antibody, S2P6, binds. There is a structural change around the D1146 residue when transitioning from the pre-fusion to the post-fusion form of the S2 region.

## Highlights

- Anti-SARS-CoV-2 bispecific Abs can be generated by combining non-neutralizing Abs
- Anti-S2 antibody CvMab-62 recognizes a novel epitope at S2 stem helix
- Bispecific antibodies can inhibit spike-mediated membrane fusogenic mechanism
- Bispecific Abs restore antiviral activity against bebtelovimab-resistant BQ.1.1

## KEY RESOURCES TABLE

| REAGENT or RESOURCE | SOURCE | IDENTIFIER |
| --- | --- | --- |
| **Antibodies** |  |  |
| CvMab-6 | Yamamoto et al.<sup>83</sup> | N/A |
| CvMab-62 | This study | N/A |
| Bebtelovimab (LY-CoV1404) | Westendorf et al.<sup>29</sup> | N/A |
| Bis1 | This study | N/A |
| Bis2 | This study | N/A |
| Bis3 | This study | N/A |
| Bis4 | This study | N/A |
| Bis-Bebtelovimab | This study | N/A |
| SARS-CoV-2 anti-spike (clone 1A9) | GeneTex Inc. | Cat# GTX632604 |
| Anti-coronavirus spike S2 antibody | Sino Biological Inc. | Cat# 40590-T62 |
| Cleaved SARS-CoV-2 S (Ser 686 ) Antibody | Cell Signaling Technology | Cat# 84534 |
| Anti-GAPDH (clone 3H12) | Medical & Biological Laboratories Co., Ltd | Cat# 171-3 |
| Horse anti-mouse IgG-HRP | Cell Signaling Technology | Cat# 7076 |
| Goat anti-mouse IgG-HRP | Proteintech Group | Cat# SA00001-1 |
| Anti-rabbit IgG-HRP | Cytiva™ | Cat# NA934 |
| Anti-His-tag mAb-HRP-Direct | MBL | Cat# D291-7 |
| Goat anti-mouse IgG (H+L) cross-adsorbed with Alexa Fluor® 488 | Thermo Fisher Scientific | Cat# A-11001 |
| **Bacterial and virus strains** |  |  |
| SARS-CoV-2 wuhan | 2019-nCoV/Japan/TY/WK-521/2020 | GISAID ID#<br>EPI_ISL_408667 |
| SARS-CoV-2 Alpha B.1.1.7 | hCoV-19/Japan/QHN001/2020 | GISAID ID#<br>EPI_ISL_804007 |
| SARS-CoV-2 Delta B.1.617.2 | hCoV-19/Japan/TY11-927/2021 | GISAID ID#<br>EPI_ISL_2158617 |
| SARS-CoV-2 Omicron BA.1 | hCoV-19/Japan/TY38-873/2021 | GISAID ID#<br>EPI_ISL_7418017 |
| SARS-CoV-2 Omicron BA.5.2.1 | hCoV-19/Japan/TY41-704/2022 | GISAID ID#<br>EPI_ISL_13241868 |
| SARS-CoV-2 BQ.1 | hCoV-19/Japan/TY41-796/2022 | GISAID ID#<br>EPI_ISL_15579783 |
| **Chemicals, peptides, and recombinant proteins** |  |  |
| Trimeric SARS-CoV-2 Spike wuhan | Sino Biological Inc. | Cat# 40589-V08H4 |
| Trimeric SARS-CoV-2 Spike BA.1 | Sino Biological Inc. | Cat# 40589-V08H26 |
| Trimeric SARS-CoV-2 Spike BA.4/5 | Sino Biological Inc. | Cat# 40589-V08H32 |
| Trimeric SARS-CoV-2 Spike BQ.1 | Sino Biological Inc. | Cat# 40589-V08H41 |
| RBD of the SARS-CoV-2 Spike wuhan | Murae et al.<sup>84</sup> | N/A |
| RBD of the SARS-CoV-2 Spike BA.1 | Murae et al.<sup>84</sup> | N/A |
| hACE2-Fc | Murae et al.<sup>84</sup> | N/A |
| His-S1 spike protein | Ray Biotech | Cat# 230-01102 |
| His-S2 spike protein | Ray Biotech | Cat# 230-01103 |
| PBS | FUJIFILM Wako Pure Chemical Corp. | Cat# 166-23555 |
| PEIpro® transfection reagent | Polyplus Transfection | Cat# 101000017 |
| Kanamycin | FUJIFILM Wako Pure Chemical Corp. | Cat# 117-00961 |
| G418 | FUJIFILM Wako Pure Chemical Corp. | Cat# 074-06801 |
| Bovine Serum Albumin fatty acid free | FUJIFILM Wako Pure Chemical Corp. | Cat# 011-15144 |
| 4%-Paraformaldehyde Phosphate Buffer Solution | NACALAI TESQUE, INC. | Cat# 09154-14 |
| 10% Tween 20 | Bio-Rad Laboratories Inc. | Cat# 1610781 |
| Sulfuric acid | FUJIFILM Wako Pure Chemical Corp. | Cat# 195-04706 |
| 1-Step™ TMB ELISA Substrate Solutions | Thermo Fisher Scientific | Cat# 34028 |
| DMEM | FUJIFILM Wako Pure Chemical Corp. | Cat# 044-29765 |
| Opti-MEM™ | Thermo Fisher Scientific | Cat# 31985062 |
| Vectashield Vibrance Antifade Mounting Medium with DAPI | Vector Laboratories Inc. | Cat# H-1700 |
| PicaGene Meliora Star-LT Luminescence Reagent | TOYO B-NET Co., Ltd. | Cat# MLT100 |
| EzWestLumi plus® | ATTO Corp. | Cat# WES-7120 |
| **Experimental models: Cell lines** |  |  |
| HEK 293T cells | ATCC | CRL-3216 |
| HEK293/ACE2 cells | This study | N/A |
| VeroE6/TMPRSS2 cells | JCRB | JCRB1819 |
| **Recombinant DNA** |  |  |
| pcDNA3.1-spike/Wuhan | Yamamoto et al. <sup>83</sup> | N/A |
| pcDNA3.1-spike/D614G | Yamamoto et al. <sup>83</sup> | N/A |
| pcDNA3.1-spike/Alpha | Yamamoto et al. <sup>83</sup> | N/A |
| pcDNA3.1-spike/Delta | Yamamoto et al. <sup>83</sup> | N/A |
| pcDNA3.1-spike/BA.1 | Yamamoto et al. <sup>83</sup> | N/A |
| pcDNA3.1-spike/BA.5.2 | This study | GenBank:<br>UPN16705.1 |
| pcDNA3.1-spike/BA.2.75 | This study | GenBank:<br>USV68346.1 |
| pcDNA3.1-spike/BA.5.2-K444T | This study | N/A |
| pcDNA3.1-spike/RaTG-13 | This study | GenBank: MN996532 |
| pcDNA3.1-spike/Khosta-2 | This study | GenBank: MZ190138 |
| pcDNA3.1-C-DYK-ACE2 | GenScript Japan Inc. | Cat# MC_0101086 |
| pcDNA3.1-C-DYK-TMPRSS2 | This study | OHu13675D,<br>GenScript Japan Inc. |
| pEGFP-C1 | BD Biotech Clontech | N/A |
| pQCXIP-GFP1-10 | Kodaka et al. <sup>52</sup> | Addgene 68715 |
| pQCXIP-BSR-GFP11 | Kodaka et al. <sup>52</sup> | Addgene 68716 |
| pTG-Luc126 | Rafique et al. <sup>85</sup> | N/A |
| phCMV-Gag-Pol 5349 | Rafique et al. <sup>85</sup> | N/A |
| **Software and algorithms** |  |  |
| GraphPad Prism 9 | GraphPad Software, Inc. | N/A |
| UCSF Chimera | Pettersen et al. <sup>86</sup> | N/A |
| ImageJ | National Institutes of Health | http://imagej.nih.gov/ij/ |
| Jalview | Waterhouse et al. <sup>87</sup> | N/A |
| Adobe® Photoshop CS4 Extended software | Adobe Systems Inc. | N/A |
| **Other** |  |  |
| TC plate 12well | SARSTEDT | Cat# 83.3921 |
| IsoPlate-96 TC white | ParkinElmer | Cat# 6005078 |
| Costar® 96well EIA/RIA Plate | Corning | Cat# 3590 |
| 4well slide chamber | Corning | Cat# 354114 |
| Sensor Chip CM5 | Cytiva | #BR100399 |
| amine coupling kit | Cytiva | #BR100050 |
| Envision 2105 | PerkinElmer | N/A |
| SpectraMax iD3 | MOLECULAR DEVICE | N/A |
| BZ-800 | KEYENCE | N/A |
| ImageQuant LAS4000 mini-image analyzer | GE Healthcare Japan Corp. | N/A |

## Notes

### Competing Interest Statement

The authors have declared no competing interest.

